# Physiological, Behavioral, and Genetic Factors that Shape Interactions in a Plant-Growth-Promoting Maize Rhizosphere Synthetic Community

**DOI:** 10.64898/2026.07.09.737579

**Authors:** Ashley A. Paulsen, Marissa N. Roghair Stroud, Larry J. Halverson

**Author notes:** Address correspondence to Larry J. Halverson. Co-first authors (Ashley A. Paulsen, Marissa N. Roghair Stroud). Order of names was determined by a coin toss.

## Abstract

Profiling microbiomes is an important way to understand the function and composition of communities in the wild, but natural microbiomes are often highly complex and often unamendable to experimentation to reveal cause and effect relationships. By using a small group of cultivable strains to represent those found in the wild, synthetic communities are one solution to this problem. Here we describe the <u>MA</u>ize <u>R</u>hizosphere <u>S</u>ynthetic <u>C</u>ommunity (MARSc), a genome-enabled 31-member bacterial community representative of the diversity found on the roots of maize grown in Iowa soils. This community is built around *Pseudomonas putida* KT2440, a model maize rhizosphere colonist and synthetic biology chassis. We characterized microbe-microbe interactions and biofilm formation of MARSc members in a variety of environmental contexts, finding that both behaviors are broadly controlled by nutrient levels. Genomic analysis and microbiome profiling of these organisms revealed that annotated biofilm genes (such as surface attachment and exopolysaccharide production) correlated to rhizosphere colonization, but neither trait correlated to *in vitro* biofilm formation. *In vitro* interactions assay findings were surprisingly consistent with co-correlations of rhizosphere abundance amongst MARSc members on roots. Finally, we found that when applied to the roots, MARSc can increase maize growth under nitrogen-limiting conditions. Altogether, MARSc is a useful tool for identifying some of the factors influencing rhizosphere microbiome assembly and will be a strong foundation for further work in this area.

**Importance:** The microbiome surrounding the roots of plants can play an integral role in plant health and growth, and it is composed of thousands of species of microbes that are specific to the plant and environment it is grown in. However, due to this complexity, little is known about the means by which microbiomes form. In this study, we developed the MAize Rhizosphere Synthetic community (MARSc) to gain a better understanding of the formation and function of microbiomes on the roots of plants. This consortium consists of 30 bacterial isolates plus *Pseudomonas putida* KT2440, a model maize root colonist. In this study, we investigated the interactions between these organisms, their genomes, cultural characteristics, and growth on the roots of maize. We show that MARSc increases maize growth under low nitrogen fertilizer, suggesting it will be a useful tool for identifying the mechanisms behind microbiome formation and plant growth promotion.

## Introduction

The microbiome is known to have vast effects on the environment around it. In plants, this community can be manipulated in ways that address current issues, including alleviating abiotic stresses such as drought (1–4) and heat (5, 6), protection from disease (7–9), or the need to decrease synthetic nitrogen inputs (10, 11). Some microbes can even help increase the food supply via plant growth promotion (12–14). The rhizosphere microbiome hosts many interactions between soil, microbes, and plants, making the establishment of causal relationships difficult and hindering our ability to predictably alter its composition (15, 16).

Both cooperative and competitive interactions may alter community structure, frequently relying on secreted public goods. Quorum sensing molecules like autoinducer-2 (AI-2) can regulate biofilm production, and siderophores such as pyoverdine can control iron competition (17, 18). Bacteria can produce phytohormones such as auxins (19), and plants can respond by exuding metabolites through their roots (20). Up to 40% of photosynthates are secreted into the rhizosphere, and their composition can significantly affect microbial metabolism (21–23). Benzoxazinoids are highly abundant in maize root exudates and act as an anti-microbial for some and a chemoattractant for others, such as plant growth-promoting *Pseudomonas putida* (24–26). The predominant secreted maize benzoxazinoid is DIMBOA (2,4-dihydroxy-7-methoxy-1,4-benzoxazin-3-one), which rapidly breaks down into MBOA (6-methoxybenzoxazolin-2-one) that can linger in the soil for months (27), thereby having a strong influence on microbial community structure (24, 28). Stressed plants can also alter root exudation patterns to recruit specific microbes to facilitate desirable interactions (29). Collectively, diffusible metabolites shape the rhizosphere microbiome in a multitude of ways, including how bacteria interact with one another.

Most bacteria are believed to live in multispecies biofilms (30), which may be influenced by one another’s secreted metabolites. Given the benefits of the biofilm lifestyle, including protection from environmental stressors and enhanced access to nutrient-rich root surfaces, the ability to form biofilms likely contributes to successful rhizosphere colonization (31–38). Biofilm formation is context-dependent, such that the type and availability of nutrients could influence cooperative and competitive behaviors. Several studies have shown root exudates have stimulated biofilm formation by multiple genera, including *Bacillus* and *Pseudomonas* (39–43).

One approach to understanding the complex rhizosphere interactions is to use a synthetic community (SynCom), comprised microbes isolated from the plant roots (44). SynComs have been used to identify keystone species in microbiomes and in biofilms, discover emergent behaviors, and untangle biofilm formation processes and rhizosphere competence traits (9, 45–47). Moreover, SynComs can link genomic information with microbial behaviors, improving our ability to predict phenotype from genotype. When coupled with plant colonization data, this may reveal traits required for successful colonization.

Here we describe the Maize Rhizosphere Synthetic community (MARSc) as a model for the maize rhizosphere microbiome. MARSc was designed around *Pseudomonas putida* KT2440 (proposed to be re-named *P. alloputida* (48)), a model rhizosphere colonist and tractable synthetic biology chassis organism (49–52). We hypothesized that bacteria isolated from the roots of maize grown in a long-term cropping system with low inorganic nitrogen inputs are more likely to possess traits related to the alleviation of low nitrogen stress than bacteria isolated from conventionally grown maize. By exploring microbe-microbe interactions, measuring biofilm formation, and mining the genomes of MARSc members, we provide insight into traits predictive of root colonization and competitiveness in the rhizosphere of maize grown under varying nitrogen fertilizer inputs. Importantly, MARSc increased maize growth under nitrogen-limiting conditions. Altogether, MARSc has enabled us to identify a variety of factors influencing rhizosphere microbiome assembly, laying the foundation for further work in this area.

## Results

### Development of the <u>MA</u>ize Rhizosphere <u>S</u>ynthetic <u>c</u>ommunity (MARSc)

We isolated bacteria from the soil, rhizosphere, and endosphere of maize grown in soil that received low inorganic fertilizer inputs as part of a long-term cropping system study (53). More than 400 isolates were classified by 16S rRNA sequencing, from which we selected 40 isolates that represented the taxonomic diversity previously reported for maize rhizospheres in the same soil (54, 55). We screened these isolates for their interactions with each other and *Pseudomonas putida* KT2440, by measuring zones of inhibition or stimulation of growth, where the spot is the “producer” and the lawn is the “receiver” (Figure 1B). We examined these interactions on a variety of nutrient rich (King’s B [KB]; ½-strength Trypticase soy agar [TSA]) or more nutrient limited (R2A) conditions, with or without FeCl_3_. While each medium had unique interactions (Figures 1A, S1), R2A had slightly less inhibitory and more stimulatory interactions than the richer ½-TSA, and 74% of interactions were consistent between the two (Figure 1C). When comparing organisms that grew on all 3 media, the prevalence and magnitude of inhibition was greater on high-nutrient KB and lowest on low-nutrient R2A. Through these assays, we eliminated phenotypically redundant isolates, selecting 31 members for MARSc, representing phylogenetically diverse genera (Figure 2, Table S1) (56).

**Figure 1.**
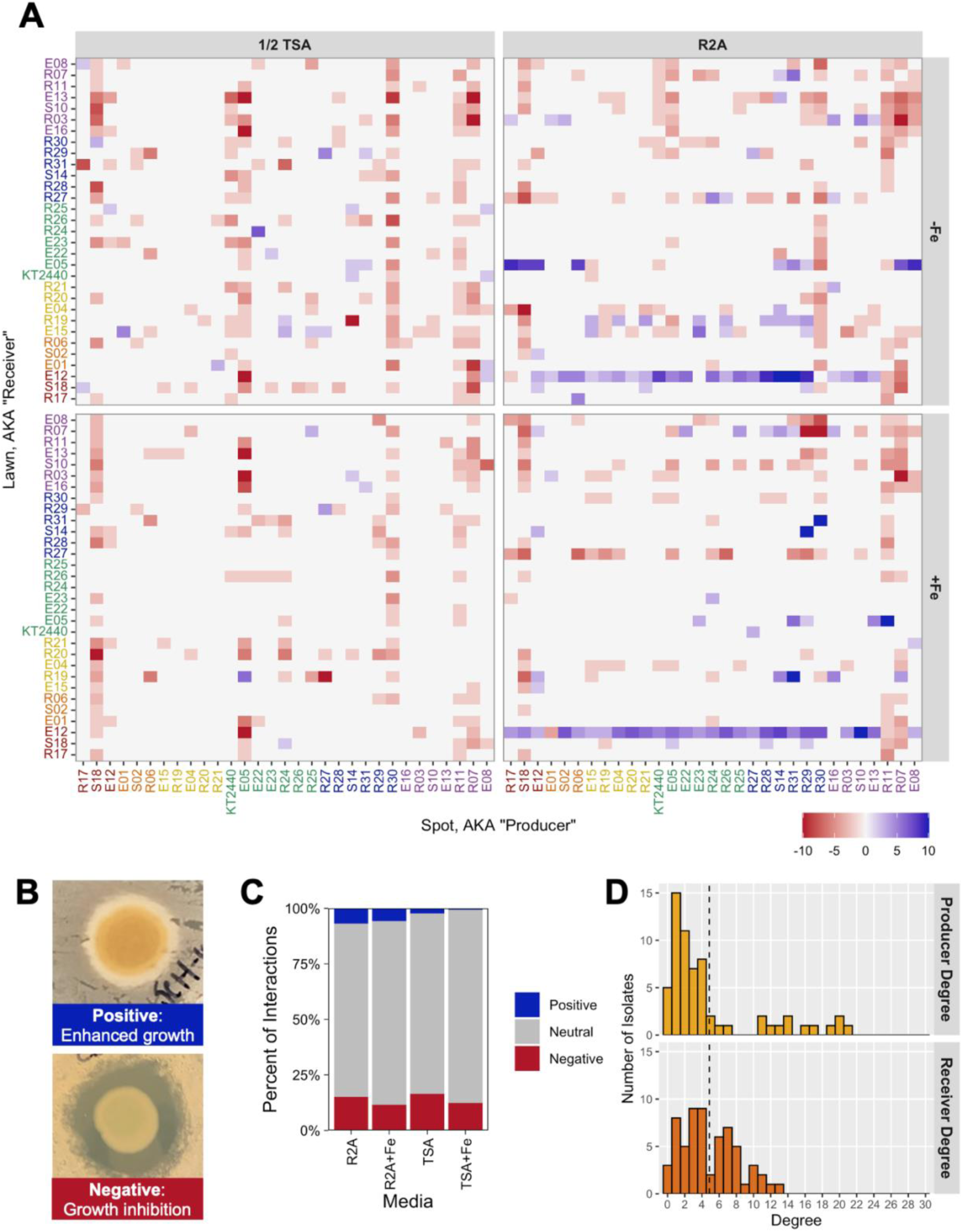
MARSc interaction assays. (A) Heat map of interactions on ½-TSA and R2A with and without supplemental iron. Blue: growth stimulation; Red: growth inhibition. (B) Illustration of positive and negative interactions. (C) Distribution of interaction types for each medium. (D) Producer and Receiver degrees for inhibitory interactions on R2A and ½-TSA without supplemental iron. Producer degree is calculated as the number of organisms the producer inhibits, while receiver degree is the number of organisms the receiver is inhibited by. Dotted line indicates the average number of inhibitory interactions as a producer or receiver.

**Figure 2.**
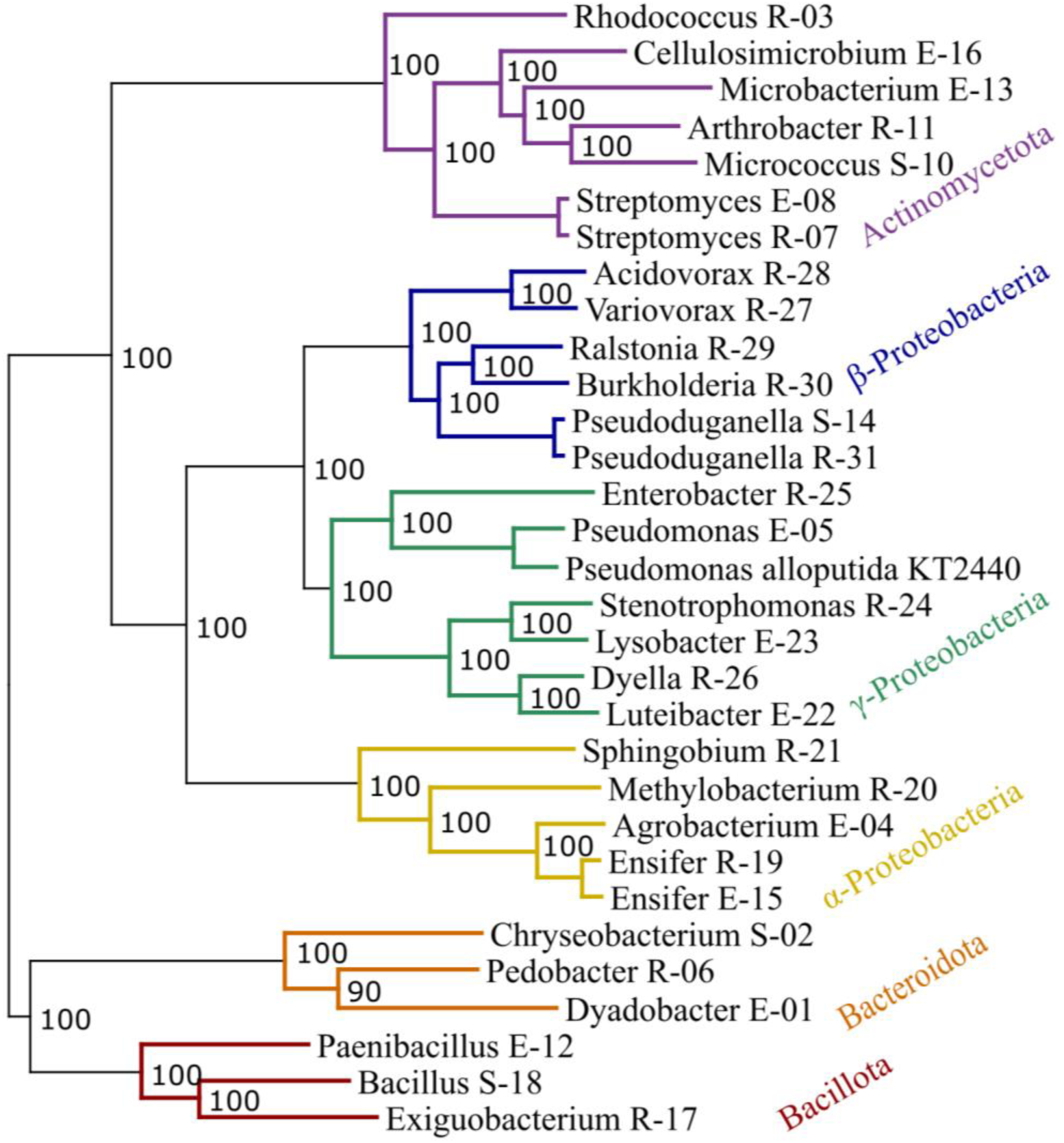
MARSc conserved gene phylogenetic tree. RAxML Maximum Likelihood Fast Bootstrapping tree with 149 single-copy gene orthologs was created using the published MARSc genomes (56) and the BV-BRC (59). Strains are identified by their genus and strain ID; strain IDs are defined by where they were obtained (E = Endosphere, R = Root, S = Soil). Phyla are color-coded as indicated in this plot. More information about MARSc members can be found in Table S1.

We further characterized phenotypes that might influence plant-microbe or microbe-microbe interactions, including production of AI-2, alkaline proteases, and indole acetic acid (IAA), as well as IAA degradation (Tables S2-S3). Autoinducer-2 production was rare while extracellular protease production was common, with varying levels of production. Six MARSc members make IAA, and four *Betaproteobacteria* can degrade it, including *Variovorax* R-27, a genus shown to reverse the inhibition of root growth promotion caused by many other bacteria (47).

### Interactions can lead to emergence of inhibitory, stimulatory, or surface colonial expansion phenotypes

In MARSc, inhibition was producer-determined while growth stimulation was receiver-determined. The leftward skewed and somewhat bimodal distribution of producer degree compared to the unimodal distribution of receiver degree (Figure 1D), as well as the greater number of vertical than horizontal red lines in the heatmaps (Figure 1A, S2A), together indicate that inhibitory interactions are determined more by the production of inhibitory metabolites than by resistance to metabolites (57). Alternatively, stimulatory interactions were often receiver-driven, as is the case for *Paenibacillus* E-12 on R2A (horizontal blue lines in Figure 1A, S2B). Some members (e.g., *Bacillus* S-18, *Burkholderia* R-30, *Streptomyce*s R-07) strongly inhibit many other MARSc strains while rarely being stimulated. In contrast, growth stimulation of *Paenibacillus* E-12 is greatest during interactions with *Beta-* and *Gamma-proteobacteria*. This outcome could be the consequence of resource limitation or auxotrophy, which is relieved by production of the metabolite(s) by other members of MARSc, thereby increasing its growth.

An unexpected observation on R2A was the occurrence of a colonial expansion phenotype by some MARSc members in the presence of other organisms. Here, colonial expansion describes the increase in colony size, which may be caused by surface motility, translocation, or increased growth (Figure S3). *Bacteroidota* members, particularly *Dyadobacter* E-01, exhibited an extensive colonial expansion phenotype, most prominently in association with *Rhizobiaceae* members (E-04, R-19, and E-15), which produce copious amounts of slime. Colonial expansion decreased in the presence of exogenous iron by an average of 2.2 mm (p < 0.0001), particularly for *Bacillus velezensis* S-18, except for *Pedobacter* R-06, which increased.

### Iron availability strongly modulates inhibitory interactions

Supplemental iron relieved many of the inhibitory interactions (Figure 1, S1). On KB, many inhibitory interactions (35%) were caused by *Burkholderia* and *Pseudomonas* species and markedly decreased when supplemented with enough FeCl_3_ to eliminate visible pyoverdine production by *Pseudomonas* strains. This indicates that Pseudomonads were more competitive for iron than many MARSc members under iron-limited conditions, likely due to the production of the high affinity siderophore pyoverdine (58). We used CRISPRi to assess whether KT2440-mediated inhibitory interactions were pyoverdine-mediated, as described previously (52). We repeated the interactions with wild type KT2440, a pyoverdine knockout (Δ*pvd*), and two CRISPRi strains (one repressing *pvdH* and one control) (Figure S4). With supplemental FeCl_3_, inhibitory interactions decreased in intensity, often disappearing completely. When the CRISPRi inducer (arabinose) was added to the medium, CRISPRi *pvdH*’s pyoverdine production was turned off to levels comparable to a KT2440 Δ*pvd* mutant, verifying inhibitory interactions by KT2440 were primarily pyoverdine-mediated. This demonstrates the importance of iron in interactions between *Pseudomonas* species and other microbes and how nutrients shape inhibitory interactions.

### Influence of root exudates and soil extracts on growth and biofilm formation

Since rhizosphere microbes live at the interface between the root and the soil, we explored how R2A media supplemented with soil extracts (SE) and maize root exudates (RE) influenced growth and biofilm forming capabilities. RE significantly stimulated growth of *Agrobacterium* E-04, *Pedobacter* R-06, and *Exiguobacterium* R-17, while growth of *Ralstonia* R-29 was inhibited. SE stimulated the growth of *Micrococcus* S-10 and *Pedobacter* R-06 and inhibited growth of *Variovorax* R-27 and *Luteibacter* E-22 (Figure 3A, Figure S5). SE had a limited effect on biofilm formation, while RE was very stimulatory (Figure 3B), though increased biofilm formation may reflect the stimulatory effect of RE on growth in general. However, some strains, including *Bacillus* S-18, only form biofilm in the presence of RE, despite no significant growth promotion by RE. Moreover, biofilm formation in response to the amendments frequently occurred differently for two strains of the same genus (*Streptomyces* E-08 and R-07; *Pseudoduganella* S-14 and R-31; *Pseudomonas* KT2440 and E-05). There were no strains that were only stimulated by SE. Collectively, RE had a greater influence on growth and biofilm formation, possibly through nutrients or signals that promote biofilm formation.

**Figure 3.**
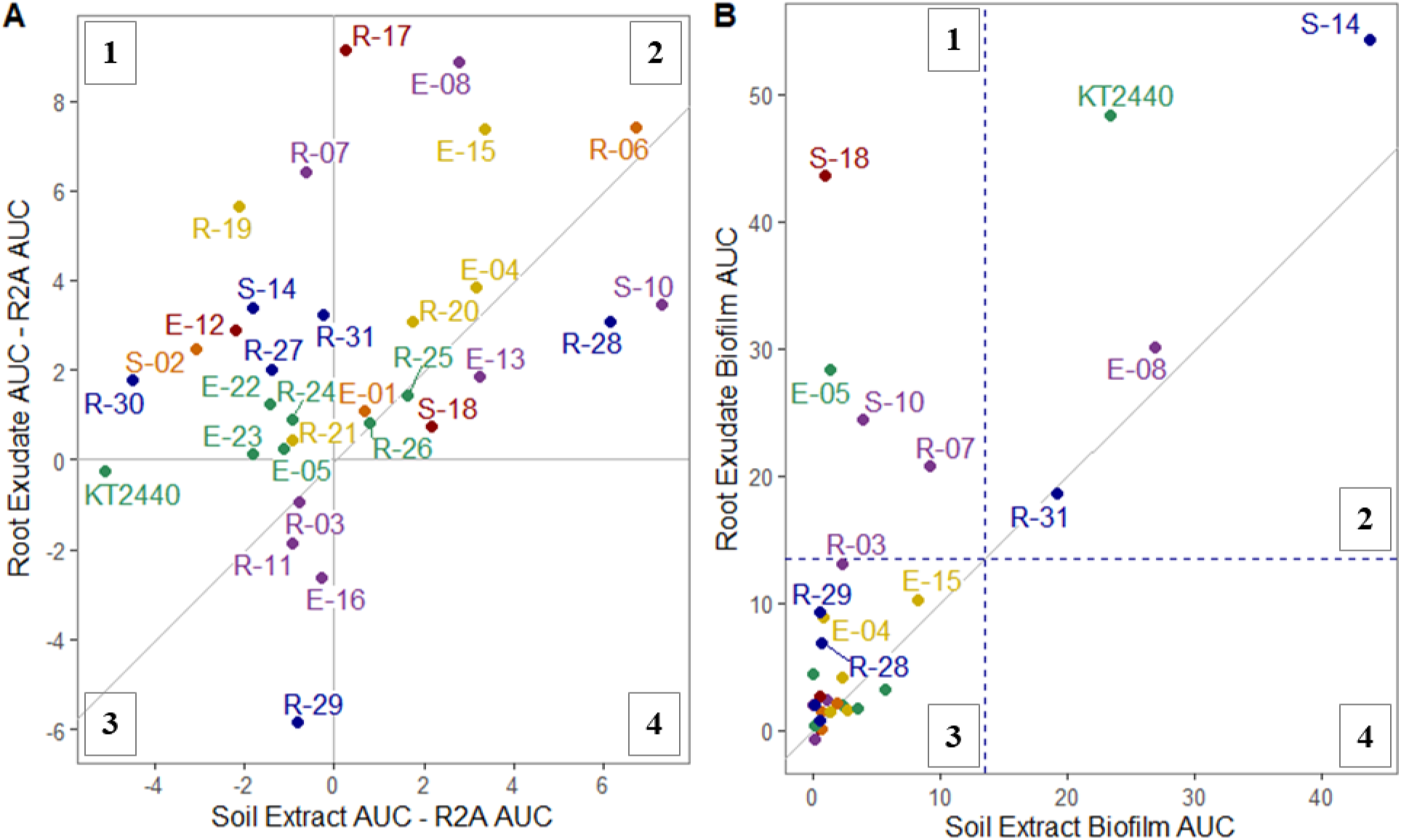
Influence of maize root exudates (RE) and aqueous soil extracts (SE) on growth and biofilm formation by MARSc members. (A) Area under the curve (AUC) of growth curves over 44 h for individual MARSc members in R2A media supplemented with RE or SE relative to the AUC in unamended R2A. The diagonal line represents similar growth in both conditions. (B) AUC of biofilm formation over 96 h in the presence of RE or SE. The dashed lines represent the cut-off for significant (p < 0.05) biofilm formation per ANOVA with Dunnett’s test against the control. Points are color-coded by phylogeny as described in Figure 2. Quadrants are labeled 1 through 4 to identify outcomes. (1) stimulated by RE and inhibited by SE, (2) stimulated by both RE and SE, (3) inhibited by both RE and SE, (4) simulated by SE and inhibited by RE.

### Effects of MBOA on growth and biofilm formation

One major component of young maize root exudates are benzoxazinoids, which are known to shape the maize rhizosphere microbiome (24, 28, 60). Of these, MBOA is the most prevalent in both concentration and persistence in the rhizosphere (27) and the most inhibitory to bacteria (24). We examined the influence of MBOA on growth and biofilm formation using a tolerance index (24), which reflects growth or biofilm formation across a range of MBOA concentrations, compared to a control. For planktonic cultures, MBOA inhibited the growth of most *Pseudomonadota* isolates, while other phyla were unaffected, though it stimulated growth of *Microbacterium* E-13 (Figure 4A). Tolerance to MBOA has been partly linked to cell wall structure (24), which may explain why the *Actinomycetota* and *Bacillota* are fairly tolerant. The significant growth inhibition of *Ralstonia* R-29 by MBOA may be a contributing factor to the growth inhibition observed in RE (Figure 3).

**Figure 4.**
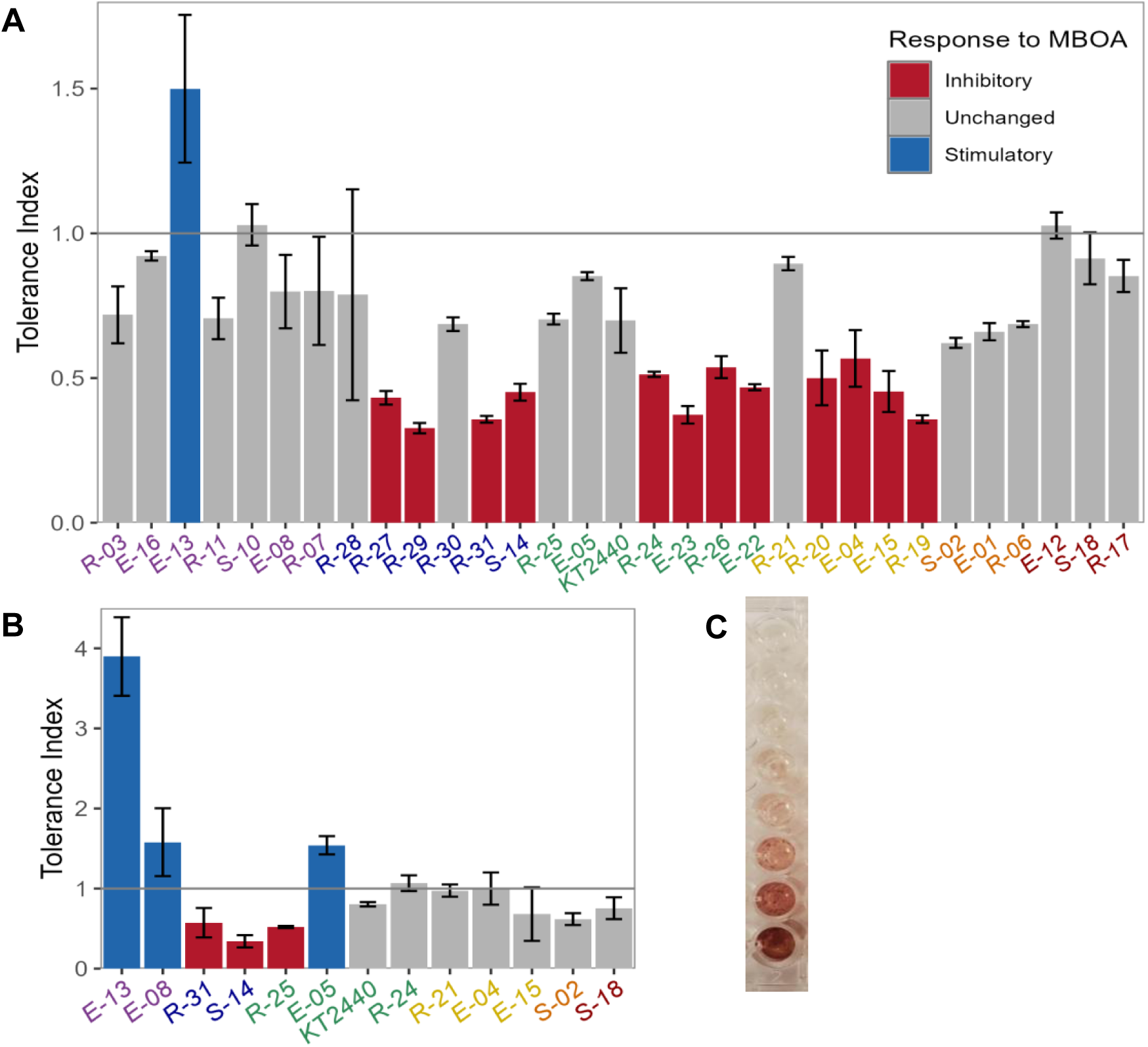
Influence of MBOA on growth and biofilm formation by MARSc members. (A) MBOA tolerance indices of planktonic cultures. The bars reflect the mean ± standard error of 3 replicates. The horizontal line represents no change in growth at the concentrations tested. Red and blue bars are significantly (p < 0.05) different from controls, based on an ANOVA with Dunnet’s post-hoc test. (B) Influence of MBOA on biofilm formation. Only strains that formed significant biofilm are shown. (C) Growth of *Microbacterium* E-13 in R2A with MBOA concentrations varying from 0 (top well) to 5000 µM (bottom well).

MBOA influenced biofilm formation differently from planktonic growth/tolerance. In R2A, 13 strains formed biofilm, and MBOA amendments inhibited biofilm formation in 3, stimulated 3, and the rest were unaffected (Figure 4B). Several *Pseudomonadota* strains inhibited by MBOA in planktonic cultures were unaffected during biofilm formation, indicating biofilm lifestyle may facilitate MBOA tolerance. Intriguingly, *Microbacterium* E-13 did not form biofilm in any other condition tested but produced a striking dark red biofilm that dramatically increased with increasing MBOA concentrations (Figure 4C). Many maize rhizosphere competent colonists, such as *Microbacterium* species, can degrade MBOA to the red-pigmented AMPO (2-amino-7-methoxyphenoxazin-3-one) by the lactonase BxdA (24, 61). BLASTP for BxdA in the *Microbacterium* E-13 genome (56) identified a lactonase capable of converting MBOA to AMPO. Yet, while many maize rhizosphere *Sphingobium* species isolates are strong AMPO-formers (24, 61), *Sphingobium* R-21 does not appear to be capable of MBOA degradation despite possessing 3 similar (62-65% identity) BxdA orthologs. Despite this, *Sphingobium* R-21 was not inhibited by MBOA, suggesting that while not visibly forming AMPO, R-21 is likely MBOA tolerant like other *Sphingobium* species. Collectively, our findings indicate that the biofilm lifestyle may influence MBOA tolerance, which could in turn influence maize rhizosphere competence.

### Influence of MARSc secreted metabolites on biofilm formation

We were interested in assessing if biofilm formation was influenced by metabolites secreted by MARSc members. Our approach was to monitor biofilm formation in a filter plate system (62, 63) that physically separates individual strains while permitting metabolites to pass through filters in the base of each well. Exposure to MARSc secreted metabolites influenced biofilm formation of 7 strains in R2A and 6 strains in R2A+RE (Figure 5A-B). These changes were not due to increased media volumes in the reservoirs, based on assays of individual strains in plates with similar volumes.

**Figure 5.**
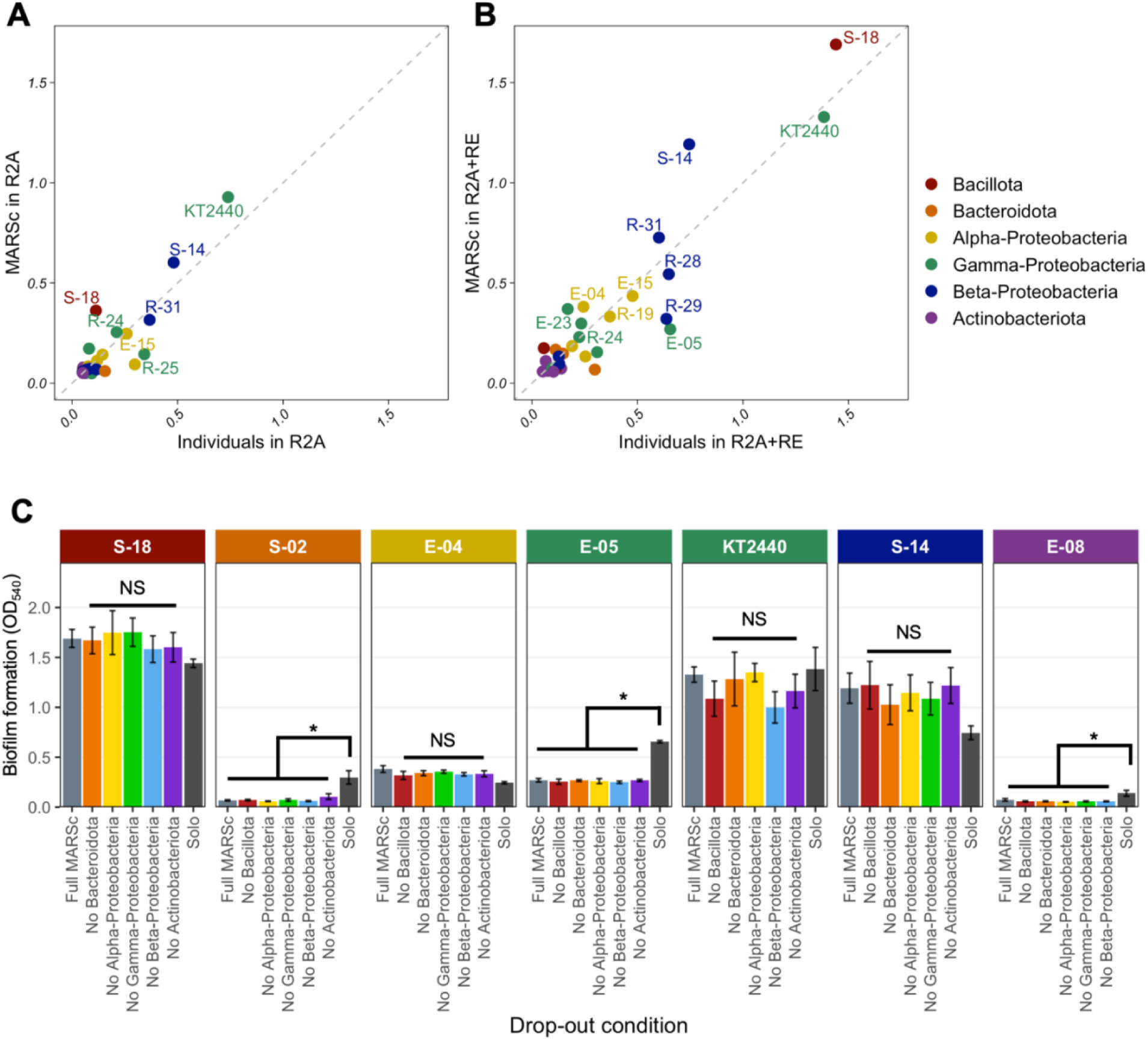
Influence of MARSc secreted metabolites on biofilm formation. Biofilm in (A) R2A and (B) R2A+RE media using a filter plate system. The y-axis is biofilm formation by individual MARSc members in the filter plate system while the x-axis is biofilm formation in normal microtiter plates. Dotted line represents when biofilm formation is equal in both conditions. (C) Influence of phylum or class dropouts on biofilm formation by select strains in R2A+RE. Bars are the mean ± standard error of 3-9 replications. Student’s t-test was used to determine significant differences in biofilm: asterisks indicate a significant difference (p < 0.05), NS: not significant.

Next, we performed drop-out assays to determine whether specific taxa were responsible for altering biofilm formation. Surprisingly, dropping out each phylum or *Pseudomonadota* class had no statistically significant effect (Figure 5C). This suggests that biofilm enhancement was not due to production of a specific metabolite(s), but instead due to depletion of inhibitory compounds by other organisms. When examining biofilm formation of individual MARSc members in media amended with spent media containing community metabolites from the filter plate system, biofilm formation in spent media only sometimes paralleled the filter plate system (Table S2). When biofilm formation was reduced in the filter plate system, it was often also reduced with spent media, indicating production of inhibitory compounds. In contrast, when biofilm formation was enhanced in the filter plate system, most organisms did not have increased biofilm formation in spent media. This indicated that in most cases, active metabolite exchange is required for stimulation and not for inhibition of biofilm formation.

### Correlations between annotated genes and biofilm formation

We were curious whether PLant-associated BActeria (PLaBAse) (64) gene categories could be used to predict biofilm formation by MARSc strains. We found no correlation between the number of annotated genes related to biofilm formation and *in vitro* biofilm formation (Figure 6A), nor was any one gene or group of genes predictive of biofilm formation. For example, *Pseudoduganella* S-14 was consistently one of the strongest biofilm formers yet had significantly fewer biofilm-related genes than *Pseudoduganella* R-31, which formed less biofilm than S-14 (Figure 6A). Moreover, there was no correlation between PLaBAse gene categories and biofilm formation or MBOA biofilm tolerance, although there were several strong primarily negative correlations with planktonic MBOA tolerance. Interestingly, the phytohormone gibberellin and GABA production gene categories had significant positive correlations to both biofilm and MBOA tolerance (Figure 6B). GABA is also associated with regulating the switch between planktonic and biofilm lifestyles in some Pseudomonads (65), which could explain positive correlations with biofilm formation. The strong correlation between gibberellin production genes and MBOA tolerance is likely due to a single ferredoxin gene found in 9 MARSc members, of which only *Methylobacterium* R-20 was significantly inhibited by MBOA (Figure 4A).

**Figure 6.**
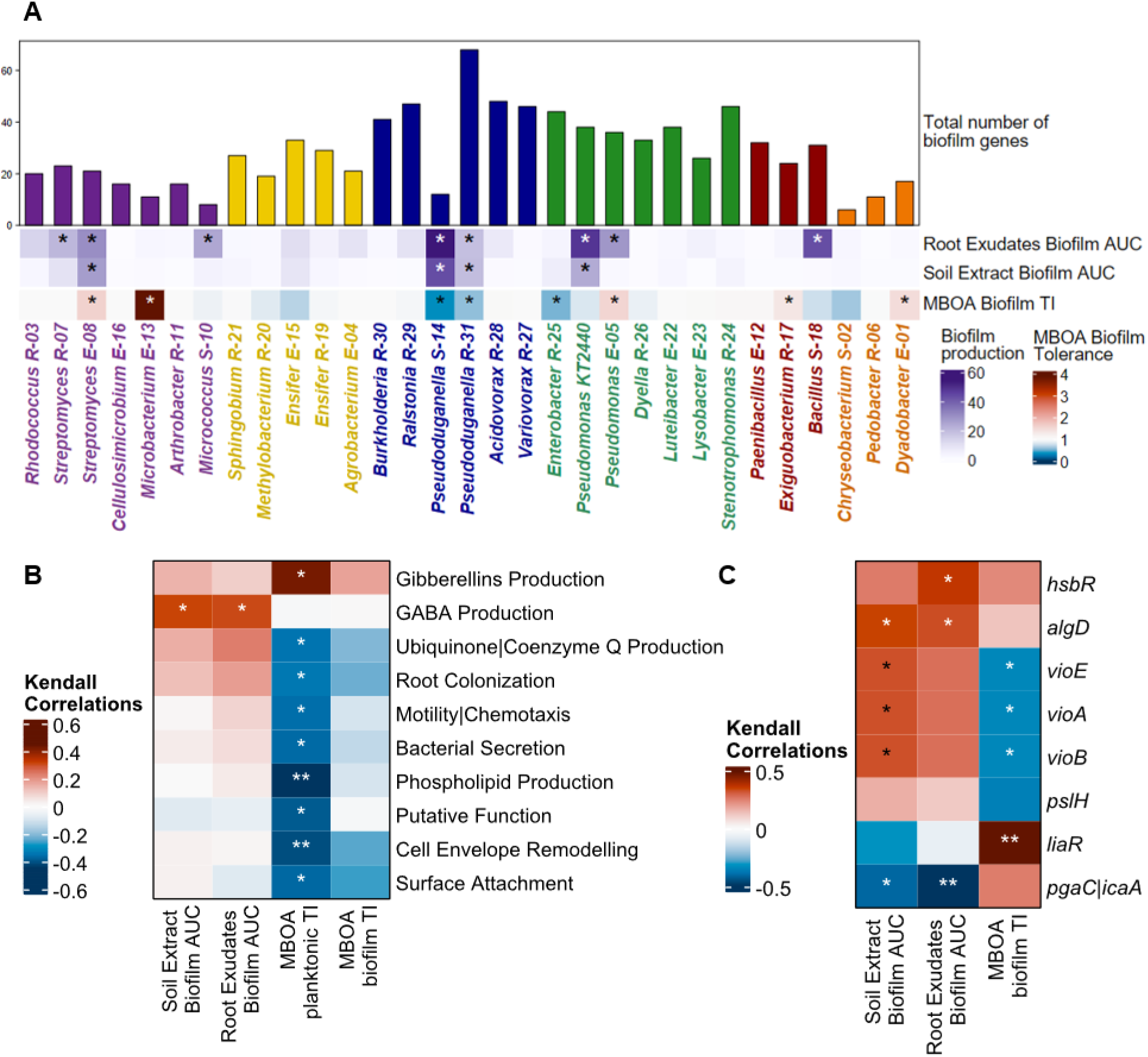
Comparison of predicted biofilm genes to biofilm phenotype. (A) Number of biofilm-associated genes (bar chart) as annotated by PLaBAse categories and biofilm formation (heat map) in R2A with RE/SE. MBOA biofilm tolerance index (TI) is defined as biofilm formation across a range of MBOA concentrations compared to a control. Asterisks denote significant (p < 0.05) biofilm formation based on a Dunnet’s test against the control. Bars and taxa color-coded by phylogeny. (B) Correlation between biofilm formation and PLaBAse categories. (C) Correlations between biofilm formation and specific biofilm-associated genes. Asterisks represent significant (p-value < 0.05) correlations. AUC = Area Under the Curve, TI = Tolerance Index.

Other genes correlated to biofilm formation include polysaccharide production (*algD*, *pgaC*|*icaA*) and a cyclic-di-GMP regulator (*hsbR*). Polysaccharide production is a vital element of biofilm formation and the HsbR-HsbA system has been shown to regulate the switch between sessile and planktonic lifestyles in *P. aeruginosa* (37, 66). Seven MARSc members possess at least one of the two. Curiously, *pgaC*|*icaA* is negatively correlated to biofilm formation, despite its role in PGA polysaccharide production (67). Importantly, *liaR* is positively correlated with biofilm formation in the presence of MBOA (Figure 6C); LiaR regulates cell envelope stress response and contributes to antibiotic resistance and biofilm formation (68–70). *liaR* orthologs are present in 22 members of MARSc, including all *Actinomycetota* and *Bacillota*. Notably, E-13, E-08, and E-05 (Figure 4B) all possess *liaR* orthologs, which suggests that the ability to adjust to cytoplasmic membrane-perturbing compounds is an important trait for biofilm formation in the presence of MBOA. The negative correlation between violacein production genes *vioABE* and biofilm formation in MBOA is likely due to the strong inhibition of MBOA on the growth of *Pseudoduganella*, the only MARSc members with *vioABE* (Figure 4B).

### Maize growth promotion and rhizosphere colonization

Previously, we observed that an earlier iteration of MARSc comprised of 40 strains exhibited biofertilizer potential when maize was grown in a soil:sand (1:1 volume) mixture fertilized at low nitrogen levels (71). We explored this further by inoculating maize seedlings with MARSc and growing them in a sterile Turface-based mix at low, moderate, or high nitrogen-fertilization rates. After 34 days, shoot biomass was significantly greater in the MARSc-inoculated plants than the untreated control for the moderate and high nitrogen levels (Figure 7). Notably, MARSc-inoculated plants with moderate fertilization had comparable biomass to the untreated controls with high nitrogen fertilization levels, highlighting its biofertilizer potential.

**Figure 7.**
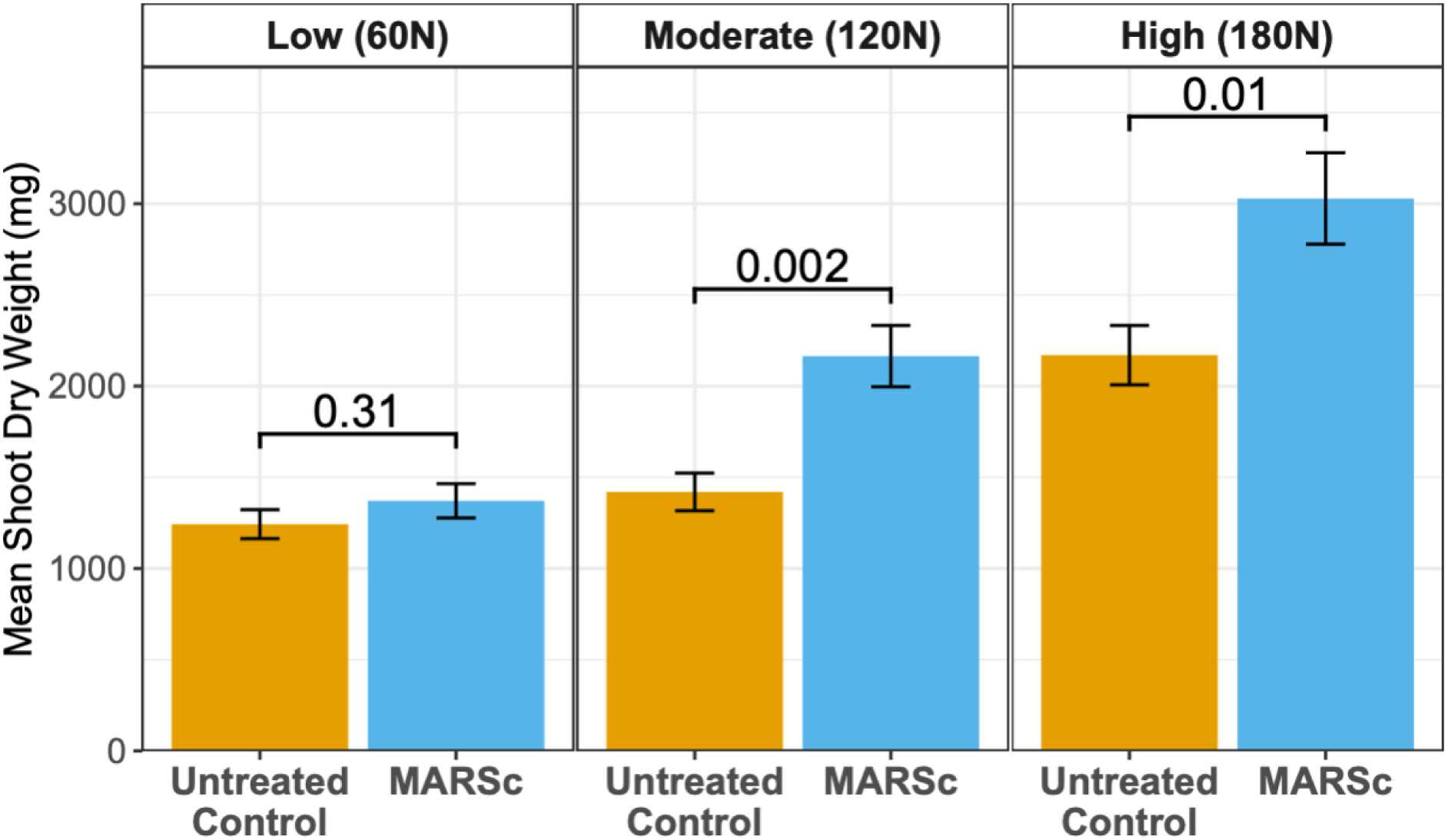
Mean shoot dry weight of untreated control and MARSc-inoculated plants grown at different nitrogen-fertilization levels (pounds of NH_4_NO_3_ nitrogen per acre). Error bars show the standard error of the mean. The brackets and values shown are the p-values from pairwise t-tests.

This finding led us to explore how nitrogen fertilization rate influences rhizosphere colonization of MARSc members 21-days post-inoculation, a time before we observed a plant growth promotion response. We used full-length 16S rRNA amplicon sequencing of the rhizosphere community to resolve MARSc members to the species level (Figure 8A). Roughly 50% of the community was derived from MARSc, with non-MARSc bacteria likely introduced either as seed endophytes, through aerial deposition, or during watering with non-sterile water. MARSc OTUs were also present in very low abundance in the untreated controls, but were significantly (ALDEx2, p < 0.05) more abundant in treated plants. The relative abundance of MARSc members in the inoculum had no apparent effect on rhizosphere colonization. Nitrogen fertilization rate also had very few effects; only *Dyadobacter* E-01 showed significant changes in abundance, decreasing as nitrogen increased (ALDEx2 with Kruskall-Wallis, p-value = 0.01). Co-correlations of MARSc members are dominated by positive correlations, except for *P. putida* KT2440 and *Variovorax* R-27, which were both consistently negatively correlated with other MARSc members (Figure 8B).

**Figure 8.**
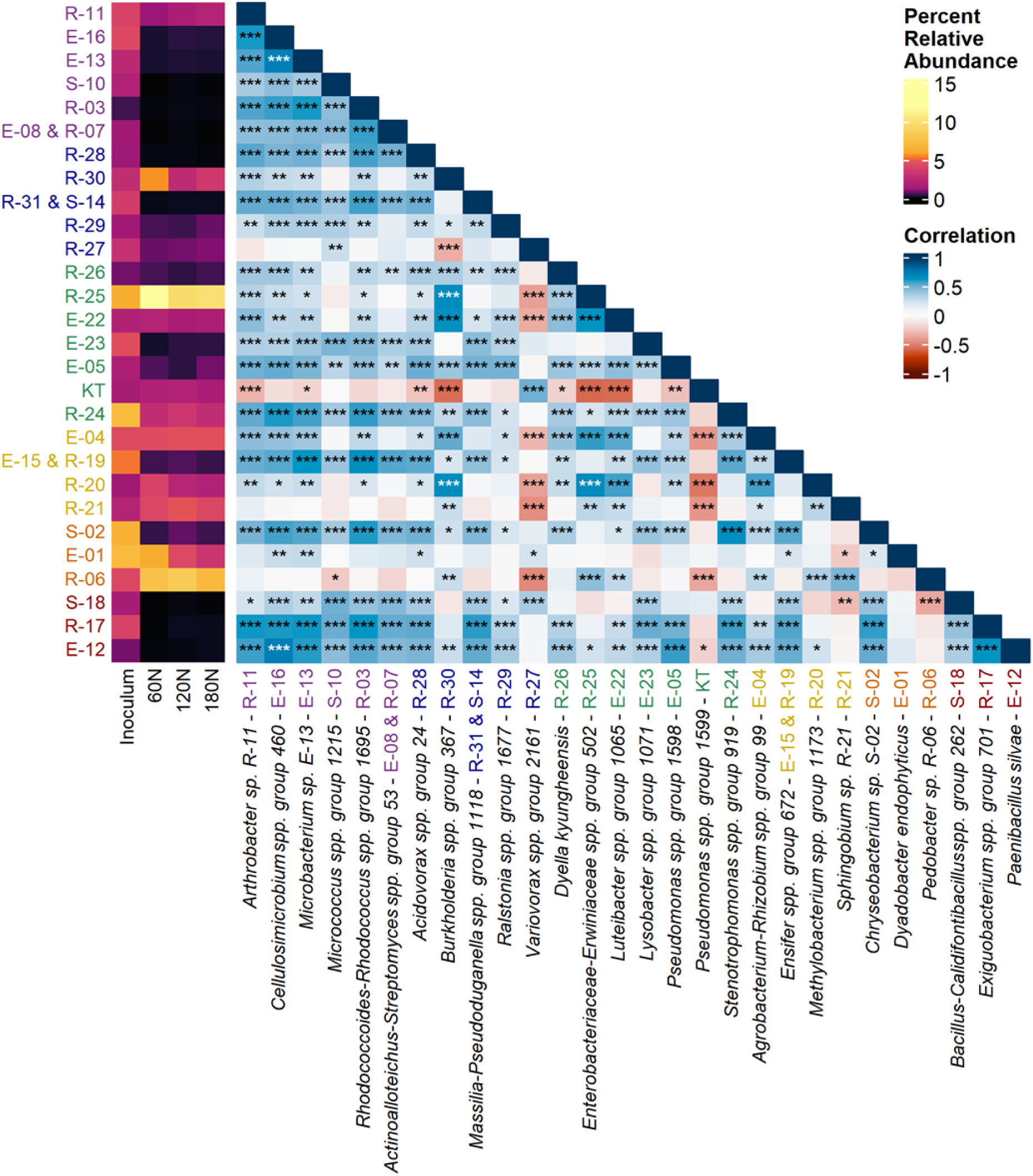
MARSc rhizosphere colonization and correlations between members. (A) Relative abundance of MARSc members in the starting inoculum compared to their relative abundance in the rhizosphere of maize grown in Turface potting mix at various nitrogen fertilization rates 21-days post-inoculation. Boxes next to each species contain the MARSc strain ID and are color coded by phyla. Relative abundance is averaged across all replicates (n=20) from two independent experiments. The minimum relative abundance detected in these samples was 0.0008%. (B) Co-correlation of relative abundance of MARSc species. The relevant species/species group for each member is paired with the corresponding MARSc ID and color coded by phyla/class. Heatmap is colored according to the Kendall correlation and asterisks indicate p-value, *** p < 0.001, **p< 0.01, *p < 0.05.

### Correlations between genotype, growth characteristics, and rhizosphere colonization

Unlike many prior studies showing biofilm formation is associated with rhizosphere competence (72–74), we did not observe this, even when RE was utilized in our biofilm assays (Figure 6). Additionally, while we did not observe any significant correlations between biofilm tolerance to MBOA and rhizosphere abundance, there was a significant negative correlation (p < 0.04) between planktonic MBOA tolerance and rhizosphere abundance. Many of the *Pseudomonadota* strains that were inhibited by MBOA had higher abundances in the rhizosphere (Figure 4A, 8) than those tolerant to MBOA, which was unexpected given how MBOA tolerance reportedly shapes the maize rhizosphere microbiome (24). There were also no significant correlations between rhizosphere abundance and IAA production/degradation or protease activity.

We looked for genes that may predict rhizosphere abundance by correlating the annotated gene content of MARSc genomes (56) with their rhizosphere abundance. COG functional categories were not correlated with rhizosphere abundance, presumably because COG categories are too broad. Similarly, there were no significant correlations to the higher level gene categories identified by the PLaBAse Plant Growth-Promoting Traits Prediction tool “PGPT-Pred” (64), despite the categories “plant-derived substrate usage” and “neutralizing abiotic stress” representing an average of 20% and 13%, respectively, of coding sequences in each member’s genome. However, we did find correlations with more precise PLaBAse categories; exopolysaccharide (EPS) production was significantly (p-value < 0.05) positively correlated to rhizosphere abundance, most notably the sub-categories of colanic acid metabolism and the Wza-Wzb-Wzc system. At an individual gene level, 7 out of the 13 significantly positively correlated genes are categorized under plant-derived substrate usage (Table S4). The 6 negatively correlated genes were much more diverse in function, without a clear pattern (Table S4).

## Discussion

Genome-enabled SynComs are important tools for revealing mechanisms behind microbiome assembly and function, including emergent behaviors that can arise within communities. In this study, we created MARSc to represent the observed phylogeny of bacteria on maize roots at the developmental stage in which maize is poised for high nitrogen uptake, with the rationale that plant selection for a microbiome facilitating nitrogen acquisition would be in place by then (54, 55). MARSc was centered around the model rhizosphere colonist and synthetic biology chassis *P. putida* KT2440, providing biotechnological flexibility. We assessed a range of physiological, behavioral, and genetic factors to not only create the SynCom but also for predicting interactions amongst members and their influence on plant health. MARSc members are rhizosphere competent and, importantly, MARSc exhibits nitrogen biofertilizer capabilities. Since plant growth promotion is observed in a synthetic growth medium, one or more MARSc members likely directly influences plant nitrogen acquisition. Yet, rhizosphere competency and plant growth promoting properties were not fully apparent from *in vitro* assays or genome mining.

*In vitro* interactions among members of MARSc were broadly controlled by nutrient levels, which is consistent with the stress-gradient hypothesis: oligotrophic conditions favor neutral or mutualistic interactions, while copiotrophic conditions favor competition (75, 76). Inhibitory interactions were generally “producer-determined”, likely resulting from the production of inhibitory compounds by the producer, rather than the presence of resistance mechanisms in the receivers (45, 57). Further, the frequency and magnitude of many inhibitory interactions were defined by iron availability. Notably, inhibition by Pseudomonads was due to production of pyoverdines, making iron less available for others and inhibiting their growth. In contrast, stimulatory interactions were “receiver-determined”, likely arising from nutrient limitations that are relieved by nearby organisms producing a limiting metabolite, such as with *Paenibacillus* E-12. Stimulation of colonial expansion/motility was an unexpected interaction between some MARSc members. Most prominently, *Dyadobacter* E-01 moved great distances, in a swarming-like pattern, when spotted atop other organisms, likely due to their production of surfactants, EPS, and/or signal molecules. As described elsewhere (45) (77, 78), this emergent behavior only occurred in low nutrient media and involved a *Bacteroidota*. Ecological advantages for this cooperativity are unclear, but it may facilitate nutrient acquisition.

The importance of diffusible metabolites was evident in the strong influence of root exudates, soil extract, and the root metabolite MBOA on growth and biofilm formation properties, although those traits were not predictive rhizosphere competence. Root exudates had more of an effect than soil extracts on planktonic growth, and stimulation of planktonic growth did not reflect stimulation of biofilm formation. While other studies have shown that *in vitro* biofilm formation or MBOA tolerance can correlate with rhizosphere colonization (22, 24, 79, 80), we did not observe similar correlations, even for *Microbacterium* E-13, where biofilm formation emerges when exposed to MBOA. Surprisingly, planktonic MBOA tolerance was generally negatively correlated with rhizosphere abundance. One possibility is that individuals who are less tolerant of MBOA rely on other microbes in the community to degrade it, such as by E-13 and possibly by other MARSc members as well. Consistent with a synergistic community-level interaction was our observation that biofilm formation by individual strains exposed to metabolites produced by others in the absence of cell-to-cell contact. Phyla/class-level drop-outs of community members did not significantly change biofilm formation, suggests there is functional redundancy within our SynCom in modulating the metabolite environment. While spent media from individual MARSc members could replicate the inhibition it did not typically replicate enhancement of biofilm formation observed in filter plates. Growth in the presence of spent media can be stimulated by decreased concentrations of inhibitory compounds or by increased availability of nutrients secreted by other microbes (cross-feeding) and can be inhibited by antibacterial compounds (81, 82). The influence of synergistic interspecies cooperation and active metabolite exchange in biofilm formation is clear (83, 84), yet how it influences the rhizosphere competence of MARSc members over time is less clear.

The best predictor of rhizosphere competence was based on genome mining, where genes for EPS production and plant derived substrate usage were significantly positively correlated with MARSc member rhizosphere abundance; both traits are known to be associated with rhizosphere colonization and/or biofilm composition (37, 85, 86). While biofilm formation may be an important rhizosphere competence trait, our approach of only examining single-species biofilms *in vitro* may not adequately reflect what occurs in mixed species communities. We observed that an average of 33% of the PLaBAse defined plant-derived substrate usage genes were significantly positively correlated with rhizosphere abundance. No other gene category was represented as substantially as plant-derived substrate usage, whether there was a positive or negative correlation with colonization. This suggests that carbon substrate usage should be explored further as a means to better predict rhizosphere competency.

Excitingly, we observed that MARSc increased the shoot biomass of maize under medium/high nitrogen fertilization levels when compared to untreated controls, when planted in a gnotobiotic synthetic substrate. An earlier iteration of MARSc showed growth promotion in a soil-sand mix with a history of low inorganic nitrogen inputs (71), but that soil contained high levels of organic nitrogen and an indigenous microbiota. These findings indicate that MARSc biofertilizer capabilities is not solely due to enhanced soil nitrogen mineralization, as our synthetic substrate did not include organic matter. Alternatively, plant growth promotion could be a consequence of the ability of MARSc to manipulate phytohormone levels in the rhizosphere. Ten MARSc members can either produce or degrade the phytohormone IAA, which has been linked to growth stimulation of plants (47) and 6 of these were present in greater than 1% relative abundance in the rhizosphere. Moreover, *Pseudoduganella* R-31 and S-14 are *Oxalobacteraceae*, which can enhance maize growth under nitrogen-limiting conditions (87). It’s possible that these MARSc members stimulate root development independent of IAA production or degradation, despite their low abundance in the rhizosphere.

*In vitro* interactions assay findings were surprisingly consistent with co-correlations of rhizosphere abundance amongst MARSc members on roots. For example, the prevalence of negative interactions between *P. putida* KT2440 and other MARSc members was consistent between experiments, while the rest of MARSc is dominated by positive correlations. *Pseudomonas* E-05 exhibited inhibitory interactions in copiotrophic conditions and stimulatory interactions in oligotrophic conditions; in the rhizosphere it had primarily positive co-correlations, which may indicate the rhizosphere is an oligotrophic growth environment. *Variovorax* R-27 also had primarily negative co-correlations in the rhizosphere, which corresponds to its frequent inhibition by other MARSc members *in vitro*. Finally, *Paenibacillus* E-12’s growth was stimulated by nearly all MARSc members, which corresponds to a high number of positive co-correlations in the rhizosphere and increased growth in the presence of MARSc metabolites.

Here, we have described MARSc, a plant-growth-promoting consortium resource for untangling the complex microbe-microbe and microbe-plant interactions occurring in the maize rhizosphere. Modeled after the maize rhizosphere community of Midwest USA corn belt soils, it has the potential to elucidate biofertilizer mechanisms, an attractive biotechnology solution to reduce inorganic fertilizers inputs into these agroecosystems. With this consortium we found that interactions between members are broadly controlled by nutrient levels, and metabolite exchange amongst members can change their behavior. Moreover, *in vitro* interactions of individual members were commonly consistent with rhizosphere colonization properties, although genome mining reveled EPS production and surface attachment were also correlated to colonization. Going forward, these findings lay the foundation for building our understanding of the interactions that shape microbiome assembly and establish MARSc as a model for studying interactions in the maize rhizosphere as well as investigating mechanisms of plant-growth-promotion.

## Abbreviated Methods

An expanded version of the methods below can be found in the supplemental information.

### Isolation of MARSc members and genome annotation

Bacteria were isolated from the bulk soil, rhizosphere, or endosphere of maize grown in soil with low inorganic nitrogen inputs from a long-term cropping system study site (55). MARSc genomes (56) were annotated with Bakta (88), reCOGnizer (89), and PLaBAse (64).

### Interaction assays

We assessed all pairwise interactions between MARSc members by spreading isolates onto solid medium, then spotting with others (45). Plates were incubated at 30°C and zones of inhibition or stimulation were measured daily in millimeters.

### Single strain biofilms

Overnight cultures were diluted to OD_600_ = 0.1 in fresh medium, then 10 µL of culture was added to 180 µL of media in a microplate with peg lids (90). Plates were incubated at 27°C. Lids were removed, rinsed in deionized water to remove planktonic cells, and dried. Biofilms were stained with crystal violet, decolorized with 33% acetic acid, and quantified by OD_540_. MBOA’s effect on biofilm formation was assessed by adding various concentrations to the medium, and the data summarized using a tolerance index (24).

### MARSc interaction biofilm assays

We used Transwell 96-well plates (filter plates) as a base for the biofilm peg lids (62). MARSc members were cultivated and diluted as above and randomly assigned to wells. The plates were incubated at room temperature (∼22°C). Lids were rinsed, stained, and quantified as above. The liquid from the plate reservoir was filtered and then used in spent media biofilm assays with individual strains using a 50/50 mix of fresh and spent media. Drop-out biofilm assays were conducted the same as the Transwell assays, with omission of each phylum/*Proteobacteria* class.

### MARSc rhizosphere colonization and maize growth promotion assays

Strains were grown in R2A, diluted to OD_600_ = 0.1 in phosphate buffer (PB), and mixed to generate the inoculum. Pre-germinated Phz51 seedlings (91) were incubated in PB or inoculum for one hour. Seedlings were planted in an autoclaved Turface mix and watered with Murashige and Skoog plant growth medium with NH_4_NO_3_ (92). For the rhizosphere colonization assay, SE was also included, and plants were grown for 21 days. Plants were harvested and the majority of the loose substrate from roots was removed. Roots were placed in PB, vortexed, and sonicated to wash off rhizosphere soil. The rhizosphere washate was centrifuged and the pellet was frozen for DNA extraction. For the maize growth promotion assay, plants were grown for 34 days, then shoots were removed, dried, and weighed.

### Full length 16S amplicon profiling

Rhizosphere samples were extracted using a Qiagen PowerSoil Pro DNA extraction kit, loading plates to decrease risk of cross-contamination between adjacent wells (93). Custom primers were ordered from IDT to combine long-read-optimized 13 bp barcodes (94) and degenerate 27F (GAGTTTGATYMTGGCTCAG) and 1492R (CGGYTACCTTGTTACGACTT) 16S rRNA primers. DNA was PCR amplified, verified by gel electrophoresis, and pooled into libraries by band intensity. Samples were sequenced by SeqCenter on a GridION sequencer. Guppy was used for super-accurate basecalling.

### Microbiome analysis

The ACT pipeline was used to generate Operational Taxonomic Unit (OTU) tables (95). Absolute abundances were calculated for each sample using the abundance of the synthetic spike-ins. Relative abundance was calculated by microeco. Differential abundances were tested using ALDEx2 (96, 97) at every taxonomic level, comparing untreated vs. treated at each nitrogen level and comparing treated plants across nitrogen levels.

## Data availability

The sequences have been deposited in the Sequence Read Archive under accession no. <u>PRJNA1442200</u>. The MARSc genomes are deposited in GenBank under accession no. <u>PRJNA1228643</u>.

## Author Contributions

LJH, MNRS, and AAP conceived the study. MNRS and AAP contributed equally to the manuscript and order of authorship was determined by a coin toss. MNRS, AAP, and LJH performed experiments and analyzed data. LJH supervised the project and provided training and mentorship. MNRS and AAP wrote the first draft of the manuscript. All authors provided comments and approved the final manuscript.

## Acknowledgements

The authors would like to thank the following individuals for their contributions to this project: Dua Vang, Hannah Burkhart, Alyssa Allard, and Katie Allgaier. Phz51 maize seeds were generously provided by Thomas Lübberstedt. This research was supported by the USDA National Institute of Food and Agriculture AFRI Education and Workforce Development Predoctoral Fellowship (2022-11424), USDA National Institute of Food and Agriculture AFRI (2021-67019-34833), National Science Foundation (DRL-1814001 and DGE-1545453) grants, the Iowa Agriculture and Home Economics Experiment Station, and the U.S. Department of Energy (DOE) Office of Biological and Environmental Research, Biological Systems Division under FWP No AL-18-380-055 to the Ames Laboratory. The Ames Laboratory is operated for the DOE by Iowa State University under contract no. DE-ACo2-07CH11358.

## References

1. Armanhi JSL, De Souza RSC, Biazotti BB, Yassitepe JEDCT, Arruda P. 2021. Modulating Drought Stress Response of Maize by a Synthetic Bacterial Community. Front Microbiol 12:747541.

2. Martins SJ, Rocha GA, de Melo HC, de Castro Georg R, Ulhôa CJ, de Campos Dianese É, Oshiquiri LH, da Cunha MG, da Rocha MR, de Araújo LG, Vaz KS, Dunlap CA. 2018. Plant-associated bacteria mitigate drought stress in soybean. Environ Sci Pollut Res 25:13676–13686.

3. Fadiji AE, Santoyo G, Yadav AN, Babalola OO. 2022. Efforts towards overcoming drought stress in crops: Revisiting the mechanisms employed by plant growth-promoting bacteria. Front Microbiol 13.

4. Yang N, Nesme J, Røder HL, Li X, Zuo Z, Petersen M, Burmølle M, Sørensen SJ. 2021. Emergent bacterial community properties induce enhanced drought tolerance in Arabidopsis. npj Biofilms and Microbiomes 7.

5. Abd El-Daim IA, Bejai S, Meijer J. 2014. Improved heat stress tolerance of wheat seedlings by bacterial seed treatment. Plant Soil 379:337–350.

6. Khan MA, Asaf S, Khan AL, Jan R, Kang S-M, Kim K-M, Lee I-J. 2020. Thermotolerance effect of plant growth-promoting Bacillus cereus SA1 on soybean during heat stress. BMC Microbiol 20:175.

7. Lee S-M, Kong HG, Song GC, Ryu C-M. 2021. Disruption of Firmicutes and Actinobacteria abundance in tomato rhizosphere causes the incidence of bacterial wilt disease. The ISME Journal 15:330–347.

8. Hoff G, Arguelles Arias A, Boubsi F, Pršić J, Meyer T, Ibrahim HMM, Steels S, Luzuriaga P, Legras A, Franzil L, Lequart-Pillon M, Rayon C, Osorio V, de Pauw E, Lara Y, Deboever E, de Coninck B, Jacques P, Deleu M, Petit E, Van Wuytswinkel O, Ongena M. 2021. Surfactin Stimulated by Pectin Molecular Patterns and Root Exudates Acts as a Key Driver of the Bacillus -Plant Mutualistic Interaction. mBio 12.

9. Niu B, Paulson JN, Zheng X, Kolter R. 2017. Simplified and representative bacterial community of maize roots. Proceedings of the National Academy of Sciences of the United States of America 114:E2450–E2459.

10. Zhang J, Liu Y-X, Zhang N, Hu B, Jin T, Xu H, Qin Y, Yan P, Zhang X, Guo X, Hui J, Cao S, Wang X, Wang C, Wang H, Qu B, Fan G, Yuan L, Garrido-Oter R, Chu C, Bai Y. 2019. NRT1.1B is associated with root microbiota composition and nitrogen use in field-grown rice. Nat Biotechnol 37:676–684.

11. Zhang L, Zhang M, Huang S, Li L, Gao Q, Wang Y, Zhang S, Huang S, Yuan L, Wen Y, Liu K, Yu X, Li D, Zhang L, Xu X, Wei H, He P, Zhou W, Philippot L, Ai C. 2022. A highly conserved core bacterial microbiota with nitrogen-fixation capacity inhabits the xylem sap in maize plants. Nature Communications 13:3361.

12. Egamberdiyeva D. 2007. The effect of plant growth promoting bacteria on growth and nutrient uptake of maize in two different soils. Applied Soil Ecology 36:184–189.

13. Ajijah N, Fiodor A, Pandey AK, Rana A, Pranaw K. 2023. Plant Growth-Promoting Bacteria (PGPB) with Biofilm-Forming Ability: A Multifaceted Agent for Sustainable Agriculture. 1. Diversity 15:112.

14. Gómez-Godínez LJ, Aguirre-Noyola JL, Martínez-Romero E, Arteaga-Garibay RI, Ireta-Moreno J, Ruvalcaba-Gómez JM. 2023. A Look at Plant-Growth-Promoting Bacteria. Plants (Basel) 12:1668.

15. Poppeliers SW, Sánchez-Gil JJ, de Jonge R. 2023. Microbes to support plant health: understanding bioinoculant success in complex conditions. Current Opinion in Microbiology 73:102286.

16. Vorholt JA, Vogel C, Carlström CI, Müller DB. 2017. Establishing Causality: Opportunities of Synthetic Communities for Plant Microbiome Research. Cell Host & Microbe 22:142–155.

17. Thompson JA, Oliveira RA, Djukovic A, Ubeda C, Xavier KB. 2015. Manipulation of the quorum sensing signal AI-2 affects the antibiotic-treated gut microbiota. Cell Reports 10:1861–1871.

18. Liu Y, Dai C, Zhou Y, Qiao J, Tang B, Yu W, Zhang R, Liu Y, Lu S-E. 2021. Pyoverdines Are Essential for the Antibacterial Activity of Pseudomonas chlororaphis YL-1 under Low-Iron Conditions. Applied and Environmental Microbiology 87:1–17.

19. Ravelo-Ortega G, Raya-González J, López-Bucio J. 2023. Compounds from rhizosphere microbes that promote plant growth. Current Opinion in Plant Biology 73:102336.

20. Sasse J, Martinoia E, Northen T. 2018. Feed Your Friends: Do Plant Exudates Shape the Root Microbiome? Trends in Plant Science 23:25–41.

21. Bais HP, Weir TL, Perry LG, Gilroy S, Vivanco JM. 2006. The role of root exudates in rhizosphere interactions with plants and other organisms. Annual Review of Plant Biology 57:233–266.

22. Wang P, Lopes LD, Lopez-Guerrero MG, van Dijk K, Alvarez S, Riethoven J-J, Schachtman DP. 2022. Natural variation in root exudation of GABA and DIMBOA impacts the maize root endosphere and rhizosphere microbiomes. Journal of Experimental Botany 73:5052–5066.

23. Seitz VA, McGivern BB, Daly RA, Chaparro JM, Borton MA, Sheflin AM, Kresovich S, Shields L, Schipanski ME, Wrighton KC, Prenni JE. 2022. Variation in Root Exudate Composition Influences Soil Microbiome Membership and Function. Applied and Environmental Microbiology 88:e00226–22.

24. Thoenen L, Giroud C, Kreuzer M, Waelchli J, Gfeller V, Deslandes-Hérold G, Mateo P, Robert CAM, Ahrens CH, Rubio-Somoza I, Bruggmann R, Erb M, Schlaeppi K. 2023. Bacterial tolerance to host-exuded specialized metabolites structures the maize root microbiome. Proceedings of the National Academy of Sciences 120:e2310134120.

25. Baatsen J, Hosaka GK, Mondin M, Azevedo JL, Hungria M, Quecine MC. 2025. Benzoxazinoids stimulate chemotaxis and act as a signaling molecule in *Azospirillum brasilense* Ab-V5, while showing minor effects on *Pseudomonas protegens* Pf-5. mBio 16:e01414–25.

26. Neal AL, Ahmad S, Gordon-Weeks R, Ton J. 2012. Benzoxazinoids in Root Exudates of Maize Attract Pseudomonas putida to the Rhizosphere. PLoS ONE 7:e35498.

27. Macías FA, Oliveros-Bastidas A, Marín D, Castellano D, Simonet AM, Molinillo JMG. 2004. Degradation Studies on Benzoxazinoids. Soil Degradation Dynamics of 2,4-Dihydroxy-7-methoxy-(2H)-1,4-benzoxazin-3(4H)-one (DIMBOA) and Its Degradation Products, Phytotoxic Allelochemicals from Gramineae. J Agric Food Chem 52:6402–6413.

28. Kudjordjie EN, Sapkota R, Steffensen SK, Fomsgaard IS, Nicolaisen M. 2019. Maize synthesized benzoxazinoids affect the host associated microbiome. Microbiome 7:59.

29. Rolfe SA, Griffiths J, Ton J. 2019. Crying out for help with root exudates: adaptive mechanisms by which stressed plants assemble health-promoting soil microbiomes. Curr Opin Microbiol 49:73–82.

30. Flemming H-C, Wuertz S. 2019. Bacteria and archaea on Earth and their abundance in biofilms. Nat Rev Microbiol 17:247–260.

31. Jefferson KK. 2004. What drives bacteria to produce a biofilm? FEMS Microbiology Letters 236:163–173.

32. Heredia-Ponce Z, Gutiérrez-Barranquero JA, Purtschert-Montenegro G, Eberl L, de Vicente A, Cazorla FM. 2021. Role of extracellular matrix components in the formation of biofilms and their contribution to the biocontrol activity of Pseudomonas chlororaphis PCL1606. Environmental Microbiology 23:2086–2101.

33. Heindl JE, Wang Y, Heckel BC, Mohari B, Feirer N, Fuqua C. 2014. Mechanisms and regulation of surface interactions and biofilm formation in Agrobacterium. Front Plant Sci 5.

34. Nilsson M, Chiang W-C, Fazli M, Gjermansen M, Givskov M, Tolker-Nielsen T. 2011. Influence of putative exopolysaccharide genes on Pseudomonas putida KT2440 biofilm stability. Environmental Microbiology 13:1357–1369.

35. Timmusk S, Grantcharova N, Wagner EGH. 2005. Paenibacillus polymyxa Invades Plant Roots and Forms Biofilms. Applied and Environmental Microbiology 71:7292–7300.

36. Martínez-Gil M, Quesada JM, Ramos-González MI, Soriano MI, de Cristóbal RE, Espinosa-Urgel M. 2013. Interplay between extracellular matrix components of *Pseudomonas putida* biofilms. Research in Microbiology 164:382–389.

37. Nielsen L, Li X, Halverson LJ. 2011. Cell–cell and cell–surface interactions mediated by cellulose and a novel exopolysaccharide contribute to Pseudomonas putida biofilm formation and fitness under water-limiting conditions. Environmental Microbiology 13:1342–1356.

38. Pandin C, Le Coq D, Canette A, Aymerich S, Briandet R. 2017. Should the biofilm mode of life be taken into consideration for microbial biocontrol agents? Microbial Biotechnology 10:719–734.

39. Ghitti E, Rolli E, Vergani L, Borin S. 2024. Flavonoids influence key rhizocompetence traits for early root colonization and PCB degradation potential of Paraburkholderia xenovorans LB400. Front Plant Sci 15.

40. Xie S, Jiang L, Wu Q, Wan W, Gan Y, Zhao L, Wen J. 2022. Maize Root Exudates Recruit Bacillus amyloliquefaciens OR2-30 to Inhibit Fusarium graminearum Infection. Phytopathology® 112:1886–1893.

41. Liu Y, Feng H, Fu R, Zhang N, Du W, Shen Q, Zhang R. 2020. Induced root-secreted d-galactose functions as a chemoattractant and enhances the biofilm formation of Bacillus velezensis SQR9 in an McpA-dependent manner. Appl Microbiol Biotechnol 104:785–797.

42. Sharma M, Saleh D, Charron J-B, Jabaji S. 2020. A Crosstalk Between Brachypodium Root Exudates, Organic Acids, and Bacillus velezensis B26, a Growth Promoting Bacterium. Front Microbiol 11.

43. Jin Y, Zhu H, Luo S, Yang W, Zhang L, Li S, Jin Q, Cao Q, Sun S, Xiao M. 2019. Role of Maize Root Exudates in Promotion of Colonization of Bacillus velezensis Strain S3-1 in Rhizosphere Soil and Root Tissue. Curr Microbiol 76:855–862.

44. Marín O, González B, Poupin MJ. 2021. From Microbial Dynamics to Functionality in the Rhizosphere: A Systematic Review of the Opportunities With Synthetic Microbial Communities. Frontiers in Plant Science 12:1–12.

45. Lozano GL, Bravo JI, Garavito Diago MF, Park HB, Hurley A, Peterson SB, Stabb EV, Crawford JM, Broderick NA, Handelsman J. 2019. Introducing THOR, a model microbiome for genetic dissection of community behavior. mBio 10:1–14.

46. Sun X, Xie J, Zheng D, Xia R, Wang W, Xun W, Huang Q, Zhang R, Kovács ÁT, Xu Z, Shen Q. 2023. Metabolic interactions affect the biomass of synthetic bacterial biofilm communities. mSystems 8:e01045–23.

47. Finkel OM, Salas-González I, Castrillo G, Conway JM, Law TF, Teixeira PJPL, Wilson ED, Fitzpatrick CR, Jones CD, Dangl JL. 2020. A single bacterial genus maintains root growth in a complex microbiome. Nature 587:103–108.

48. De Lorenzo V, Pérez-Pantoja D, Ramos JL, Nikel PI. 2026. The disputed identity of *Pseudomonas putida* KT2440: when taxonomists rename your favorite microbe. mBio e03390–25.

49. Espinosa-Urgel M, Salido A, Ramos JL. 2000. Genetic analysis of functions involved in adhesion of Pseudomonas putida to seeds. J Bacteriol 182:2363–2369.

50. Ramos-González MI, Campos MJ, Ramos JL. 2005. Analysis of Pseudomonas putida KT2440 gene expression in the maize rhizosphere: in vivo [corrected] expression technology capture and identification of root-activated promoters. J Bacteriol 187:4033–4041.

51. De Lorenzo V, Pérez-Pantoja D, Nikel PI. 2024. *Pseudomonas putida* KT2440: the long journey of a soil-dweller to become a synthetic biology chassis. J Bacteriol e00136–24.

52. Roghair Stroud MN, Vang DX, Halverson LJ. 2024. Optimized CRISPR Interference System for Investigating Pseudomonas alloputida Genes Involved in Rhizosphere Microbiome Assembly. ACS Synth Biol 13:2912–2925.

53. Davis AS, Hill JD, Chase CA, Johanns AM, Liebman M. 2012. Increasing Cropping System Diversity Balances Productivity, Profitability and Environmental Health. PLOS ONE 7:e47149.

54. Wattenburger CJ, Halverson LJ, Hofmockel KS. 2019. Agricultural management affects root-associated microbiome recruitment over maize development. Phytobiomes Journal 3:260–272.

55. Bay G, Lee C, Chen C, Mahal NK, Castellano MJ, Hofmockel KS, Halverson LJ. 2021. Agricultural Management Affects the Active Rhizosphere Bacterial Community Composition and Nitrification. mSystems 6:10.1128/msystems.00651-21.

56. Paulsen AA, Vang DX, Halverson LJ. 2025. Complete genome sequences of 37 bacteria in a maize rhizosphere synthetic community. Microbiology Resource Announcements 0:e00496–25.

57. Vetsigian K, Jajoo R, Kishony R. 2011. Structure and evolution of streptomyces interaction networks in soil and in silico. PLoS Biology 9:e1001184.

58. Schalk IJ, Perraud Q. 2023. *Pseudomonas aeruginosa* and its multiple strategies to access iron. Environmental Microbiology 25:811–831.

59. Olson RD, Assaf R, Brettin T, Conrad N, Cucinell C, Davis JJ, Dempsey DM, Dickerman A, Dietrich EM, Kenyon RW, Kuscuoglu M, Lefkowitz EJ, Lu J, Machi D, Macken C, Mao C, Niewiadomska A, Nguyen M, Olsen GJ, Overbeek JC, Parrello B, Parrello V, Porter JS, Pusch GD, Shukla M, Singh I, Stewart L, Tan G, Thomas C, VanOeffelen M, Vonstein V, Wallace ZS, Warren AS, Wattam AR, Xia F, Yoo H, Zhang Y, Zmasek CM, Scheuermann RH, Stevens RL. 2023. Introducing the Bacterial and Viral Bioinformatics Resource Center (BV-BRC): a resource combining PATRIC, IRD and ViPR. Nucleic Acids Res 51:D678–D689.

60. Cotton TEA, Pétriacq P, Cameron DD, Meselmani MA, Schwarzenbacher R, Rolfe SA, Ton J. 2019. Metabolic regulation of the maize rhizobiome by benzoxazinoids. ISME J 13:1647–1658.

61. Thoenen L, Kreuzer M, Pestalozzi C, Florean M, Mateo P, Züst T, Wei A, Giroud C, Rouyer L, Gfeller V, Notter MD, Knoch E, Hapfelmeier S, Becker C, Schandry N, Robert CAM, Köllner TG, Bruggmann R, Erb M, Schlaeppi K. 2024. The lactonase BxdA mediates metabolic specialisation of maize root bacteria to benzoxazinoids. Nat Commun 15:6535.

62. Chodkowski JL, Shade A. 2017. A Synthetic Community System for Probing Microbial Interactions Driven by Exometabolites. mSystems 2:4–7.

63. Chodkowski JL, Shade A. 2024. Bioactive exometabolites drive maintenance competition in simple bacterial communities. mSystems 9:e00064–24.

64. Patz S, Gautam A, Becker M, Ruppel S, Rodríguez-Palenzuela P, Huson D. 2021. PLaBAse: A comprehensive web resource for analyzing the plant growth-promoting potential of plant-associated bacteria. bioRxiv 10.1101/2021.12.13.472471.

65. Takeuchi K. 2018. GABA, A Primary Metabolite Controlled by the Gac/Rsm Regulatory Pathway, Favors a Planktonic Over a Biofilm Lifestyle in *Pseudomonas protegens* CHA0. MPMI 31:274–282.

66. Bouillet S, Ba M, Houot L, Iobbi-Nivol C, Bordi C. 2019. Connected partner-switches control the life style of Pseudomonas aeruginosa through RpoS regulation. Sci Rep 9:6496.

67. Steiner S, Lori C, Boehm A, Jenal U. 2012. Allosteric activation of exopolysaccharide synthesis through cyclic di-GMP-stimulated protein–protein interaction. The EMBO Journal 10.1038/emboj.2012.315.

68. Klinzing DC, Ishmael N, Hotopp JCD, Tettelin H, Shields KR, Madoff LC, Puopolo KM. 2013. The two-component response regulator LiaR regulates cell wall stress responses, pili expression and virulence in group B Streptococcus. Microbiology (Reading) 159:1521–1534.

69. Suntharalingam P, Senadheera MD, Mair RW, Lévesque CM, Cvitkovitch DG. 2009. The LiaFSR System Regulates the Cell Envelope Stress Response in Streptococcus mutans. J Bacteriol 191:2973–2984.

70. Jordan S, Junker A, Helmann JD, Mascher T. 2006. Regulation of LiaRS-Dependent Gene Expression in Bacillus subtilis: Identification of Inhibitor Proteins, Regulator Binding Sites, and Target Genes of a Conserved Cell Envelope Stress-Sensing Two-Component System. Journal of Bacteriology 188:5153–5166.

71. Knight C. 2022. Does N fertilizer rate affect microbial benefits to early maize growth? An evaluation of Iowa-isolated microbial communities., 0 ed. Iowa State University, Ames (Iowa).

72. Al-Ali A, Deravel J, Krier F, Béchet M, Ongena M, Jacques P. 2018. Biofilm formation is determinant in tomato rhizosphere colonization by Bacillus velezensis FZB42. Environ Sci Pollut Res 25:29910–29920.

73. Walker TS, Bais HP, Déziel E, Schweizer HP, Rahme LG, Fall R, Vivanco JM. 2004. Pseudomonas aeruginosa-Plant Root Interactions. Pathogenicity, Biofilm Formation, and Root Exudation. Plant Physiology 134:320–331.

74. Danhorn T, Fuqua C. 2007. Biofilm Formation by Plant-Associated Bacteria. Annu Rev Microbiol 61:401–422.

75. Yan B, Liu N, Liu M, Du X, Shang F, Huang Y. 2021. Soil actinobacteria tend to have neutral interactions with other co-occurring microorganisms, especially under oligotrophic conditions. Environmental Microbiology 23:4126–4140.

76. Kawai T, Tokeshi M. 2007. Testing the facilitation–competition paradigm under the stress-gradient hypothesis: decoupling multiple stress factors. Proc R Soc B 274:2503–2508.

77. McCully LM, Graslie J, McGraw AR, Bitzer AS, Sigurbjörnsdóttir AM, Vilhelmsson O, Silby MW. 2021. Exploration of Social Spreading Reveals That This Behavior Is Prevalent among Pedobacter and Pseudomonas fluorescens Isolates and That There Are Variations in the Induction of the Phenotype. Applied and Environmental Microbiology 87:1–17.

78. McCully LM, Bitzer AS, Seaton SC, Smith LM, Silby MW. 2019. Interspecies Social Spreading: Interaction between Two Sessile Soil Bacteria Leads to Emergence of Surface Motility. mSphere 4:1–16.

79. Gastélum G, Ángeles-Morales A, Arellano-Wattenbarger G, Coronado Y, Guevara-Hernandez E, Rocha J. 2024. Biofilm formation and maize root-colonization of seed-endophytic Bacilli isolated from native maize landraces. Applied Soil Ecology 199:105390.

80. Liu Y, Gates AD, Liu Z, Duque Q, Schmidt SS, Chen MY, Hamilton CD, O’Toole GA, Haney CH. 2026. In vitro biofilm formation by a beneficial bacterium partially predicts in planta protection against rhizosphere pathogens. ISME J 20:wraf114.

81. Dos Santos AR, Martino RD, Testa S, Mitri S. 2022. Classifying interactions in a synthetic bacterial community is hindered by inhibitory growth medium. mSystems 10.1128/msystems.00239-22.

82. Murillo-Roos M, Abdullah HSM, Debbar M, Ueberschaar N, Agler MT. 2022. Cross-feeding niches among commensal leaf bacteria are shaped by the interaction of strain-level diversity and resource availability. ISME Journal 16:2280–2289.

83. Burmølle M, Hansen LH, Sørensen SJ. 2007. Establishment and Early Succession of a Multispecies Biofilm Composed of Soil Bacteria. Microb Ecol 54:352–362.

84. Madsen JS, Røder HL, Russel J, Sørensen H, Burmølle M, Sørensen SJ. 2016. Coexistence facilitates interspecific biofilm formation in complex microbial communities. Environmental Microbiology 18:2565–2574.

85. Carezzano ME, Alvarez Strazzi FB, Pérez V, Bogino P, Giordano W. 2023. Exopolysaccharides Synthesized by Rhizospheric Bacteria: A Review Focused on Their Roles in Protecting Plants against Stress. 4. Applied Microbiology 3:1249–1261.

86. Janczarek M, Rachwał K, Cieśla J, Ginalska G, Bieganowski A. 2015. Production of exopolysaccharide by Rhizobium leguminosarum bv. trifolii and its role in bacterial attachment and surface properties. Plant Soil 388:211–227.

87. Yu P, He X, Baer M, Beirinckx S, Tian T, Moya YAT, Zhang X, Deichmann M, Frey FP, Bresgen V, Li C, Razavi BS, Schaaf G, von Wirén N, Su Z, Bucher M, Tsuda K, Goormachtig S, Chen X, Hochholdinger F. 2021. Plant flavones enrich rhizosphere Oxalobacteraceae to improve maize performance under nitrogen deprivation. Nature Plants 7:481–499.

88. Schwengers O, Jelonek L, Dieckmann MA, Beyvers S, Blom J, Goesmann A. 2021. Bakta: rapid and standardized annotation of bacterial genomes via alignment-free sequence identification. Microbial Genomics 7:000685.

89. Sequeira JC, Rocha M, Alves MM, Salvador AF. 2022. UPIMAPI, reCOGnizer and KEGGCharter: Bioinformatics tools for functional annotation and visualization of (meta)-omics datasets. Computational and Structural Biotechnology Journal 20:1798–1810.

90. Ceri H, Olson ME, Stremick C, Read RR, Morck D, Buret A. 1999. The Calgary Biofilm Device: New Technology for Rapid Determination of Antibiotic Susceptibilities of Bacterial Biofilms. J Clin Microbiol 37:1771–1776.

91. Brenner EA, Blanco M, Gardner C, Lübberstedt T. 2012. Genotypic and phenotypic characterization of isogenic doubled haploid exotic introgression lines in maize. Mol Breeding 30:1001–1016.

92. Murashige T, Skoog F. 1962. A Revised Medium for Rapid Growth and Bio Assays with Tobacco Tissue Cultures. Physiol Plant 15:473–497.

93. Custer GF, Dibner RR. 2020. Modified Methods for Loading of High-Throughput DNA Extraction Plates Reduce Potential for Contamination. JoVE 61405.

94. Srivathsan A, Lee L, Katoh K, Hartop E, Kutty SN, Wong J, Yeo D, Meier R. 2021. ONTbarcoder and MinION barcodes aid biodiversity discovery and identification by everyone, for everyone. BMC Biology 19:217.

95. Paulsen AA, LaSarre B, Delp D, Beattie GA, Halverson LJ. 2026. A Bioinformatic Pipeline for Consensus Taxonomic Classification of Long-Read Amplicons. bioRxiv 10.64898/2026.04.29.721641.

96. Fernandes AD, Reid JN, Macklaim JM, McMurrough TA, Edgell DR, Gloor GB. 2014. Unifying the analysis of high-throughput sequencing datasets: characterizing RNA-seq, 16S rRNA gene sequencing and selective growth experiments by compositional data analysis. Microbiome 2:15.

97. Fernandes AD, Macklaim JM, Linn TG, Reid G, Gloor GB. 2013. ANOVA-Like Differential Expression (ALDEx) Analysis for Mixed Population RNA-Seq. PLOS ONE 8:e67019.

