## Supplemental Information for "Physiological, Behavioral, and Genetic Factors that Shape Interactions in a Plant-Growth-Promoting Maize Rhizosphere Synthetic Community"

Ashley A. Paulsen[1, 2], Marissa N. Roghair Stroud[1,2], Larry J. Halverson[1,2]
[1]Department of Plant Pathology, Entomology, and Microbiology, Iowa State University, Ames, IA, 50011
[2]Ames National Laboratory, Ames, IA, 50011

#### Methods

##### Seed sterilization and germination

Phz51 maize seeds were generously provided by Thomas Lübberstedt (1). Phz51 seeds were surface sterilized as we have previously described (2), by soaking in 70% ethanol for 10 minutes then 20% bleach for 10 minutes, washing five times with sterile water after each soak. Seeds were placed on germination paper moistened with a plant growth nutrient solution (1X MS + Captan fungicide (2 g/L)), then rolled and placed vertically in a beaker with a small amount of the nutrient solution in the bottom (3). Seedlings were incubated for 3-5 days until most had a 1-cm radicle emerging.

##### Bacterial growth media

Bacteria in this study were grown on the following media: King's B (KB), Reasoner's 2A medium (R2A), tryptic soy agar (TSA), which was used at full, half, and one-tenth strength (TSA, ½-TSA, and 1/10-TSA), 50% Nutrient agar (50% NA), Luria Broth/Agar (LB/LA), and Nitrogen-fixing PhotoHeterotroph Enrichment Medium with supplemental ammonium chloride (NPHEM, adapted from Gest et al. (4)). R2A was supplemented with aqueous soil extracts (SE) or maize root exudates (RE) in some experiments (R2A+SE or R2A+RE). Plants were watered with Murashige and Skoog plant growth medium (MS) with macro- and micro-nutrients (MSP21), supplemented with 1.65 g/L ammonium nitrate (3). Phosphate buffer (PB) (0.8 mM potassium phosphate dibasic, 0.2 mM potassium phosphate monobasic, pH 7.1-7.4) was used for dilutions and to collect bacteria from roots.

Aqueous soil extracts were generated using soil from the Marsden long-term cropping site (5). 100 g of soil were added to 1 L of water, which was heated to 80˚C and stirred for 16 h, before filtering out solids and autoclaving (2, 6). Maize root exudates were collected from hydroponically grown sterile maize seedlings in a system similar to one we have previously described (2). Seeds were sterilized and germinated as above. Syringe barrels (60 mL) were filled with 5 mm glass beads and seeds were planted 1 inch below the top of the syringe, then filled to the top with more glass beads and watered with 15 mL of 0.5X MS. To maintain sterility, syringes were covered with Whirl-Pak bags and ends were plugged with luer-lock valves and caps. Root exudates were aseptically collected at 7- and 9-days post-planting by removing the luer-lock cap from the valve, attaching a syringe, and pulling off the liquid from the base of the plant. Next, fresh watering solution was replaced using a second syringe through the luer-lock valve. At both timepoints, sterility of the exudates was monitored by spotting onto R2A plates and growing for 2-3 days at 26˚C. Exudates were stored at -80˚C. Following multiple rounds of collecting root exudates, samples were thawed, pooled, and passed through a 0.22 µm filter before aliquoting and re-freezing at -80˚C until use.

**Isolation and identification of MARSc members**

MARSc members were isolated as described in (7). Briefly, bacteria were isolated from the bulk soil, rhizosphere, or endosphere of V4-V5 maize grown in soil with a history of low inorganic nitrogen inputs from a long-term cropping system study site (8). Over 400 organisms were isolated, tentatively identified by sequencing their full 16S rRNA gene, and included in MARSc based on the interaction assay results as described below. Information about all strains included in this study can be found in Table S1. The phylogenetic tree was created using the BV-BRC (9) tool 'Bacterial Genome Tree' using the published MARSc genomes (7). It is a RAxML Maximum Likelihood Fast Bootstrapping based on 149 single-copy gene orthologs identified by the 'Bacterial Genome Tree' tool.

**Interaction assays**

Interaction assays based on those described by Lozano et al. (10) were performed to identify inhibitory and stimulatory interactions among MARSc members. In this assay, one isolate (referred to as the receiver) was spread onto a solid medium in a petri plate to create a lawn before aliquoting 10 µL of a cell suspension that had 100-fold more cells (based on absorbance at an optical density of 600 nm, $OD_{600}$) of another isolate (referred to as producer). There were four aliquots of the producer spotted on each receiver lawn, one per quadrant, and plates were incubated at 30°C. Solid media consisted of KB, R2A, and ½-TSA with and without 50 µM or 35 µM $FeCl_3$ amendments. These concentrations of iron were sufficient to completely repress visible pyoverdine production by all *Pseudomonas* isolates. Inhibitory interactions were based on measured sizes of zones of inhibition of the receiver surrounding the spots of the producer. Stimulatory interactions were based on enhancement of growth of the receiver or enhanced colonial expansion (motility) of the spots, as illustrated in Figure S2, which were measured in millimeters. Interactions 1 mm or less were removed, as they were often not reproducible between replicates. A paired T-test was used to compare colonial expansion between R2A and R2A+Fe in the interactions assay.

Not all isolates were able to grow on KB, full-strength TSA, 50% NA, or LA, likely preferring oligotrophic over copiotroph growth conditions. Given the scale of these assays, they were only performed once for each medium, although a subset of interactions was repeated to check for consistency, and 69% of interactions were consistent in type across the replicates (e.g. inhibitory in both replicates). However, 89% of interactions were within 2 mm of each other among replicates (the discrepancy arising most often from when one replicate showed a neutral interaction and the other was slightly inhibitory), indicating that this assay is fairly reproducible.

**Generation of interaction networks**

We generated networks to display our interaction data, as performed by others previously (10, 11). Interaction networks were generated using Cytoscape version 3.8.2 (12) and Gephi version 0.10.1 (13). Data tables of interactions (measured in millimeters of the zone of inhibition/stimulation) were uploaded to Cytoscape to generate a preliminary network. Cytoscape data was exported to Gephi for visual manipulation. Networks in Gephi display arrows pointing from the producer to the receiver of the interaction, with thickness of arrow depicting strength of interaction and color depicting type of interaction. Only interactions greater than 1 mm in diameter were included in these figures.

**CRISPRi interactions**

We inserted the arabinose-inducible dCas9 expression system designed by Peters et al. and the sgRNA expression vector pSpyB designed in our previous study into KT2440 using methods we have described before (2, 14). We used 4 strains of KT2440 for this assay, whose descriptions can be found in Table S5. We selected MARSc members to participate in this assay based on if they were previously inhibited by KT2440 in an iron-dependent manner on each of the three previously described types of media. All strains were grown in KB, ½-TSA, or R2A broth overnight, then all environmental isolates were diluted 1:100 in fresh media and spread onto the surface of petri plates with corresponding media to create lawns. A subset of plates contained $FeCl_3$ (50 µM for KB, 35 µM for others) or 66 mM arabinose. Once lawns were dry, each was spotted with the four KT2440 strains. Plates were incubated at 30˚C for 48 or 72 hours and imaged and measured daily under bright light and UV. Some strains, particularly on R2A but also on ½-TSA, showed greatly enhanced growth in the presence of arabinose.

**Growth Curves**

Overnight cultures of each strain were diluted to $OD_{600}$ = 0.1 +/- 0.005 in fresh medium, then 20 µL of the dilution was added to 200 µL of fresh corresponding medium in a 96-well plate, then incubated at 25˚C for 44 hours with shaking, measuring $OD_{600}$ hourly. Media tested included R2A, R2A+RE, and R2A+SE. Each strain had 3-6 technical replicates per plate and at least 3 independent biological replicates for each medium and each strain. Technical replicates were averaged to produce one growth curve per biological replicate per medium. The area under the curve was calculated for each of these growth curves. Differences among media were independently evaluated for each strain with one-way ANOVA and Tukey's HSD.

**Single strain biofilms**

Overnight cultures in R2A were diluted to $OD_{600}$ = 0.1 +/- 0.005 in fresh medium. Nunc™ Edge™ 96-Well, Non-Treated, Flat-Bottom Microplates with Nunc™ Immuno TSP Lids (peg lids) were used as a Calgary biofilm device (15). Each well was filled with 150 µL of fresh media and 10 µL of diluted culture. Each plate contained a minimum of 6 technical replicates per strain and an empty control of un-inoculated media. Reservoirs were filled with sterile water to help limit evaporation. Plates were covered with peg lids then wrapped in parafilm and foil and incubated at 27˚C for 24, 48, and 96 hours. To harvest, lids were removed and dipped three times in deionized water to rinse off planktonic cells, then left to dry. Lids were placed in new 96-well plates filled with 0.1% crystal violet (CV) and placed on a rotator for 15 minutes, then rinsed and dried again. Lids were decolorized with 33% acetic acid for 15 min before the acetic acid solution was quantified by $OD_{540}$ in a Biotek Synergy HI plate reader. At least 3 biological replicates were performed per strain. To determine the amount of biofilm formed, each replicate was normalized against its corresponding empty control prior to calculating the Area Under the Curve (AUC) after 48 h of growth. A one-way ANOVA with comparison against the control via Dunnet's method, with a p-value cut-off of 0.05, was used to determine if that strain formed significant amounts of biofilm in that condition.

**Transwell plate biofilm assays**

We used the Transwell 96-well plates (filter plates), described previously by Chodkowski and Shade (16, 17), as a base for the biofilm peg lids. For this system, we used the lids of the filter plates as a reservoir, filled the reservoir with 40 mL of media, and placed the filter plate atop the reservoir so the media could soak into the wells. Overnight cultures of MARSc members grown in R2A were diluted to an $OD_{600}$ of 0.1 +/- 0.005 in fresh R2A, then 10 µL of culture were added to appropriate wells, with at least 3 technical replicates per strain. In all cases, the location of the strains was randomly assigned. Finally, a peg lid was placed atop the filter plate. It should be noted that these systems are not inherently compatible: the notched corners of the peg lids had to be removed using sterile pliers in order for the lid to fit into the system. Due to this imperfect fit between lids and bases, the plates were left in the biosafety cabinet, at room temperature (about 22˚C) and in the dark for 48 or 96 hours. Following incubation, lids were gently removed and rinsed as above, then allowed to dry. 100 µL of the planktonic cells within the wells of the plates were transferred to fresh 96-well plates and measured at $OD_{600}$ to track how many planktonic cells were present in the wells. The liquid from the reservoir of the filter plate was collected, spotted onto a fresh R2A plate to check for sterility, passed through a 0.22 µm filter, and frozen at -80˚C for later use as spent medium. Peg lids were stained and quantified as above. To assess if biofilm formation was altered by the presence of MARSc metabolites using a Transwell plate, a two-way ANOVA was performed (Strain and MARSc metabolites were main effects), which was assessed for R2A and R2A+RE media individually, and a Student's T-test was used to identify significant differences.

**Drop-out and spent media biofilm assays**

Drop-out biofilm assays were conducted the same as the Transwell biofilm plate assays, with the exception of the community composition involved. Each of the 3 phyla and 3 *Proteobacteria* classes were removed one at a time from MARSc, leaving 6 drop-out communities. Spent media biofilm assays were conducted the same as the single strain assays, with the exception that the media was 50% fresh media and 50% spent media from the Transwell plate reservoir, collected as above. Both assays were stained and quantified as above. A two-way ANOVA was performed (Strain and spent/fresh media were main effects) to analyze if spent media could replicate the effects of MARSc metabolites. A Student's T-test was used to identify significant differences. Dropout assays for biofilm formation were analyzed individually for each strain, using a one-way ANOVA (main effect was the phyla/class removed), followed by a Student's T-test.

**MBOA tolerance index**

The effects of MBOA on growth and biofilm formation were assayed in the same manner as described above, with the addition of MBOA to the growth medium. A 500 mM MBOA stock solution in DMSO was further diluted in DMSO to prepare 10, 50, 125, 250 mM stocks. These dilutions ensured that a uniform volume of DMSO (10 µL/mL) was added to achieve each working concentration. Isolates were grown in R2A with MBOA concentrations of 0, 100, 500, 1250, 2500, and 5000 µM, with the 0 µM control receiving 10 µL/mL DMSO. The tolerance index was calculated as laid out by Thoenen et. al. (18). Briefly, the AUC was calculated at each level of treatment and plotted against MBOA concentrations. The AUC of this line was then divided by the AUC of the control across all concentrations to produce a tolerance index (TI). Significant promotion or inhibition was determined by one-way ANOVA with comparison against the control via Dunnet's method, with a p-value cut-off of 0.05.

**Maize growth promotion assay**

The MARSc inoculum was prepared by culturing each strain in R2A broth, diluting each overnight culture to $OD_{600}$ = 0.1 +/- 0.005 in 1 mM phosphate buffer, and mixing equal volumes of each dilution. Rhizotrons were prepared as described previously (8), except they were filled with 1300 g of an autoclaved Turface mix, composed of 2-parts Turface MVP®, 1-part Turface Quick Dry®, and 1-part vermiculite by volume. One day before planting they were watered with

1000 mL of a nutrient solution containing 0.5X MS supplemented with 39, 78, or 118 mg $NH_4NO_3$ to achieve the equivalent of 60, 120, or 180 pounds of fertilizer N per acre. Pre-germinated seedlings were sorted by size, evenly distributed across treatments, and incubated in sterile phosphate buffer or MARSc inoculum for 1 hour at room temperature. Seedlings were transferred to sterile petri dishes to allow excess inoculum to drip off before planting. Two seedlings were planted per rhizotron and then covered with an additional 250 g Turface mix prior to watering with 100 mL sterile water. Rhizotrons were placed randomly into a Percival growth chamber with a 16-h/8-h light/dark photoperiod at 24°C and watered as needed. Shoots were removed at the Turface-air interface and dried in a 60°C oven for 72 h. A Student's T-test was used to compare untreated to MARSc-treated shoot dry weights for each nitrogen level.

**MARSc rhizosphere colonization**

The MARSc inoculum was prepared by culturing each strain in R2A broth, diluting each overnight culture to $OD_{600}$ = 0.1 +/- 0.005 in 1 mM PB, and mixing equal volumes of each dilution. One mL aliquots of the initial inoculum were centrifuged at 10,000 RPM for 5 minutes, had their supernatant removed, and were stored at -80° C for amplicon profiling. Pre-germinated seedlings were prepared and inoculated as described above.

Seedlings were planted in cone-tainers, prepared as follows. A 2.5 cm circle of steel mesh was placed into the bottom of each cone-tainer, then cones were soaked in 70% ethanol for 15 minutes and air-dried. Sterile cones were filled with 80g of a pre-autoclaved Turface mix described above with the addition of 1-part peat by volume. Approximately 24 hours before planting, each cone was watered with 70 mL of 1X soil extract + 0.5X MS supplemented with 48 or 72 mg/cone $NH_4NO_3$, corresponding to 120 or 180 pounds of N per acre. One seedling was planted in each cone followed by a 100 µL drench of the corresponding inoculum. Cones were filled to the top with substrate and watered with 5 mL of the corresponding MS+N fertilizer treatment. There were 10 replicates per treatment for each experiment, which was repeated twice.

Plants were grown for 21 days in racks in a growth chamber with a 16-h/8-h light/dark photoperiod at 24 °C and watered as needed to maintain consistent moisture based on gravimetric water content. Rack positions were regularly rotated in the growth chamber to prevent edge effects. To harvest, plants were removed from cones and sterile forceps used to remove the majority of the loose substrate from the roots. The root was placed in a 50 mL conical tube with sterile phosphate buffer, vortexed for 10 seconds, and placed in a sonicating water bath for 3 minutes to wash rhizosphere soil from the root. The root was removed using sterile forceps and the rhizosphere washate centrifuged at 8,000 RPM for 5 minutes, the supernatant discarded, and the pellet of rhizosphere soil frozen at -80° C for later DNA extraction.

**Full length 16S amplicon profiling**

Rhizosphere samples were thawed and extracted using a Qiagen PowerSoil Pro 96-well plate DNA extraction kit. Plates were loaded using a modified method to decrease risk of cross-contamination between adjacent wells (19). Custom primers were ordered from IDT to combine ONT optimized 13 bp barcodes (20) and 27F (GAGTTTGATYMTGGCTCAG) and 1492R (CGGYTACCTTGTTACGACTT) 16S rRNA primers. To quantify absolute abundance, synthetic DNA spike-ins consisting of 4 unique sequences mixed in a ratio of 1000:100:10:1 were added to each reaction. However, due to difficulties detecting adequate reads for all 4 spike-ins, we were unable to use these as intended. DNA was PCR amplified in 50 µL reactions consisting of 5%

v/v DMSO, 1.5 µL of 10 mM dNTPs, 10 µL of 5X KAPA Fidelity buffer, 0.5 µL of KAPA HiFi HotStart Polymerase, 1 µL of 0.5 µM barcoded 27F primer, 1 µL of 0.5 µM barcoded 1492R primer, 2.5 µL of synthetic DNA spike-in, and 5 ng total community DNA using the following protocol: 95°C for 5 minutes, 25 cycles of 45 seconds at 98°C, 45 seconds at 54 °C, 100 seconds at 72°C, and a final extension of 72 °C for 5 minutes. Each reaction was verified by gel electrophoresis and pooled into libraries based on band intensity by classifying bands as either faint or bright and pooling at a 2:1 volume ratio. Pooled libraries were sent to SeqCenter and prepared and sequenced per the following methods. Sample libraries were prepared using the PCR-free Oxford Nanopore Technologies (ONT) Ligation Sequencing Kit with the NEBNext® Companion Module to manufacturer's specifications. Nanopore sequencing was performed on an Oxford Nanopore GridION sequencer using R10.4.1 flow cells in one or more multiplexed shared flow-cell runs. Run design utilized the 400bps sequencing mode with a minimum read length of 200bp. Adaptive sampling was not enabled. Guppy v6.5.7 was used for super-accurate basecalling (SUP), demultiplexing, and adapter removal (basecalling model dna_r10.4.1_e8.2_400bps_modbases_5mc_cg_sup). No quality trimming was performed during basecalling.

#### Microbiome analysis

The ACT pipeline was used to generate Operational Taxonomic Unit (OTU) tables (21). Abundance tables were analyzed using the R package microeco (22). Relative abundance was calculated by microeco. Differential abundances were tested using ALDEx2 (23, 24) at every taxonomic level, comparing untreated vs. treated at each nitrogen level and comparing treated plants across nitrogen levels. Correlations were performed using Kendall's method and p-values adjusted for False Discovery Rate (FDR), with $p < 0.05$ designated as significant. Beta-diversity was calculated within microeco, using Bray-Curtis dissimilarities and tested with PERMDISP and PERMANOVA.

Despite starting with sterilized substrate and surface-sterilized seeds, we detected non-MARSc bacteria in both treated and untreated plants. Pairwise PERMANOVA of β-diversity measured between inoculated and un-inoculated plants at each nitrogen level were significantly different ($p < 0.005$) but were not significantly different between nitrogen levels within a treatment. Here, we only present results on MARSc-inoculated plants.

#### Genome mining and comparison

MARSc genomes (7) were annotated with Bakta v1.9.3 (default settings) (25). Genomes were further annotated with reCOGnizer (26) to generate COG annotations and with the PLant-associated BActeria web resource (PLaBAse) (27) tool for the annotation of plant growth-promoting traits (PGPT-Pred) using BLASTp+HMMER. Annotations results were loaded as tables into R for analyses. The proportion of coding DNA sequences for each PLaBAse category was calculated within each strain by summing the gene counts by category and dividing by the total CDS for that strain. Total CDS was retrieved from the PGAP annotations described in Paulsen et al. (7). Biofilm genes were identified as any genes in the PLaBAse categories BIOFILM_REGULATORS or BIOFILM_RELATED and the total number calculated as the sum of the copy number for each gene. For correlations, genes were grouped by PLaBAse categories and/or COG categories, and the gene counts were summed within each category. All correlations are Kendall correlations with FDR (false discovery rate) adjusted p-values with a significance cut-off of $p < 0.05$.

#### Measuring other community- and plant-relevant traits

Individual MARSc members were also assessed for their ability to produce autoinducer-2 (AI-2), produce an alkaline protease, and produce or degrade indole acetic acid (IAA). AI-2 production was measured as previously described (28, 29). MARSc genomes were also screened for the *luxS* gene, which is responsible for AI-2 production, and the results matched with those that had measurable AI-2 production *in vitro*.

For the IAA assay, colonies were used to inoculate 1 mL R2A supplemented with either 0.1% tryptophan to measure production or 0.4 mM IAA to measure degradation. After shaking at 30° C for 72 hours, cells were pelleted (6,000 x g, 5 minutes) and the supernatant collected. The IAA concentration was measured using an adaption of the Gordon and Weber protocol (30). Briefly, 50 µL of supernatant was combined with 100 µL of Salkowski's reagent (10 mM $FeCl_3$, 97% reagent grade, and 34.3% perchloric acid, ACS grade) in a clear, flat-bottom 96-well plate, incubated at room temperature for 30 minutes, and the absorbance at 530 nm measured on a Biotek Synergy H1 plate reader. The IAA concentration was calculated using an IAA standard curve (0, 25, 50, 100, 250, 500, and 750 µM IAA in R2A).

Extracellular casein hydrolysis of individual MARSc members was examined on R2A or ½-strength TSA amended with 1.5 % skim milk powder (Difco). Cells were cultivated overnight in either R2A or ½-TSA broth, diluted to an OD600 of 0.01 in 1 mM phosphate buffer prior to placing 5 µL aliquots onto the same medium in which they were cultivated. Zones of clearing were measured after 1, 2, or 5 days. The experiment was repeated 3 times.

#### Statistical Analysis and Graphing

Data was analyzed and plots were created with R v4.4.3, and R package ComplexHeatmap v2.23.1 (31) was used for select heatmaps.

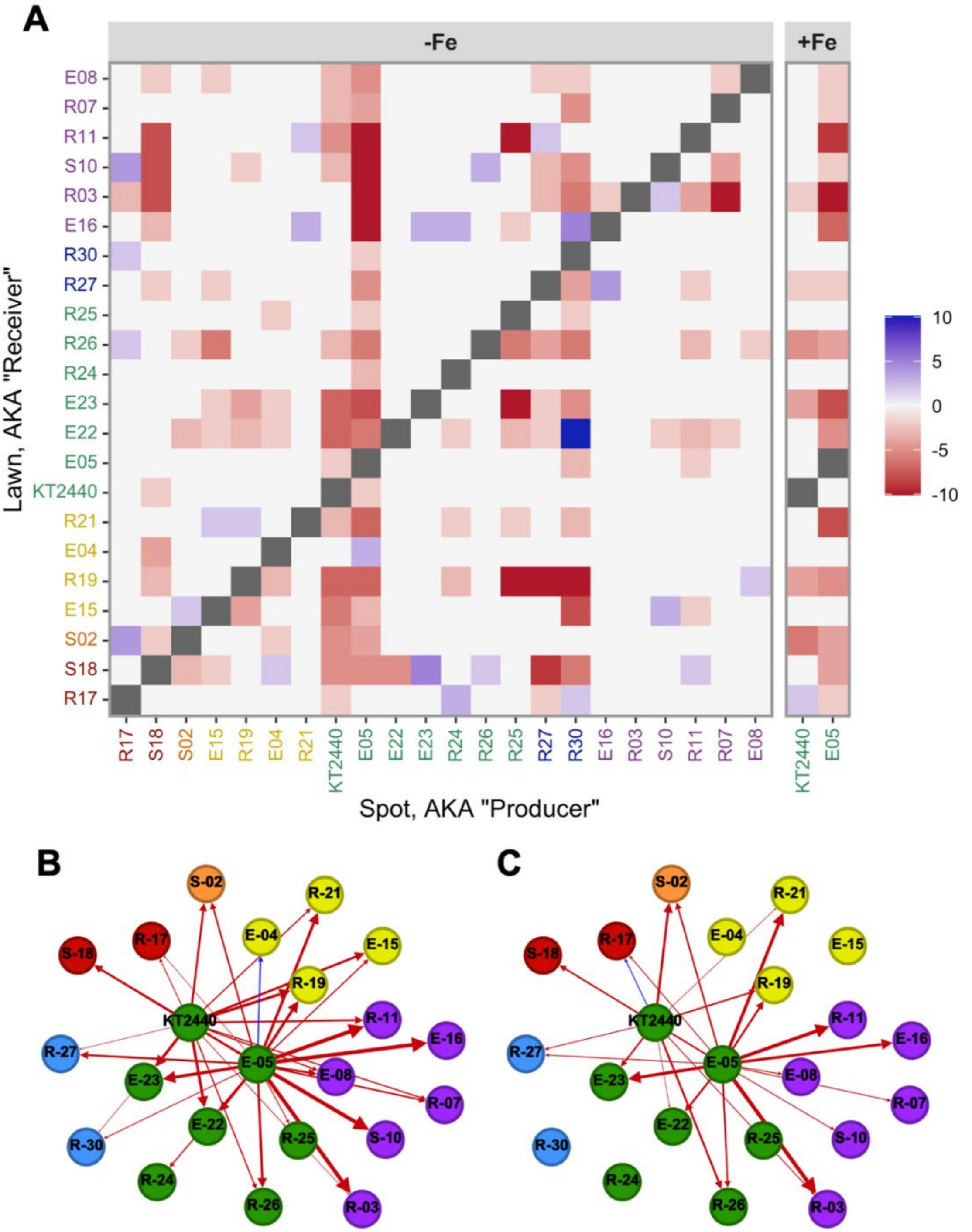


Figure S1. Interactions between isolates and *P. putida* KT2440. (A) Heat map of interactions on KB with and without 50 µM $FeCl_3$. Scale bar reflects the strength of the interaction. Blue: stimulation; Red: inhibition. Gray: not measured. The color of the strain IDs reflects their phylogeny. Red: *Bacillota*; Orange: *Bacteroidota*; Yellow: α-*Proteobacteria*; Green: γ-*Proteobacteria*; Blue: β-*Proteobacteria*; Purple: *Actinomycetota*. Interactions network with *Pseudomonas* species in the (B) absence and (C) presence of exogenous 50 µM $FeCl_3$.

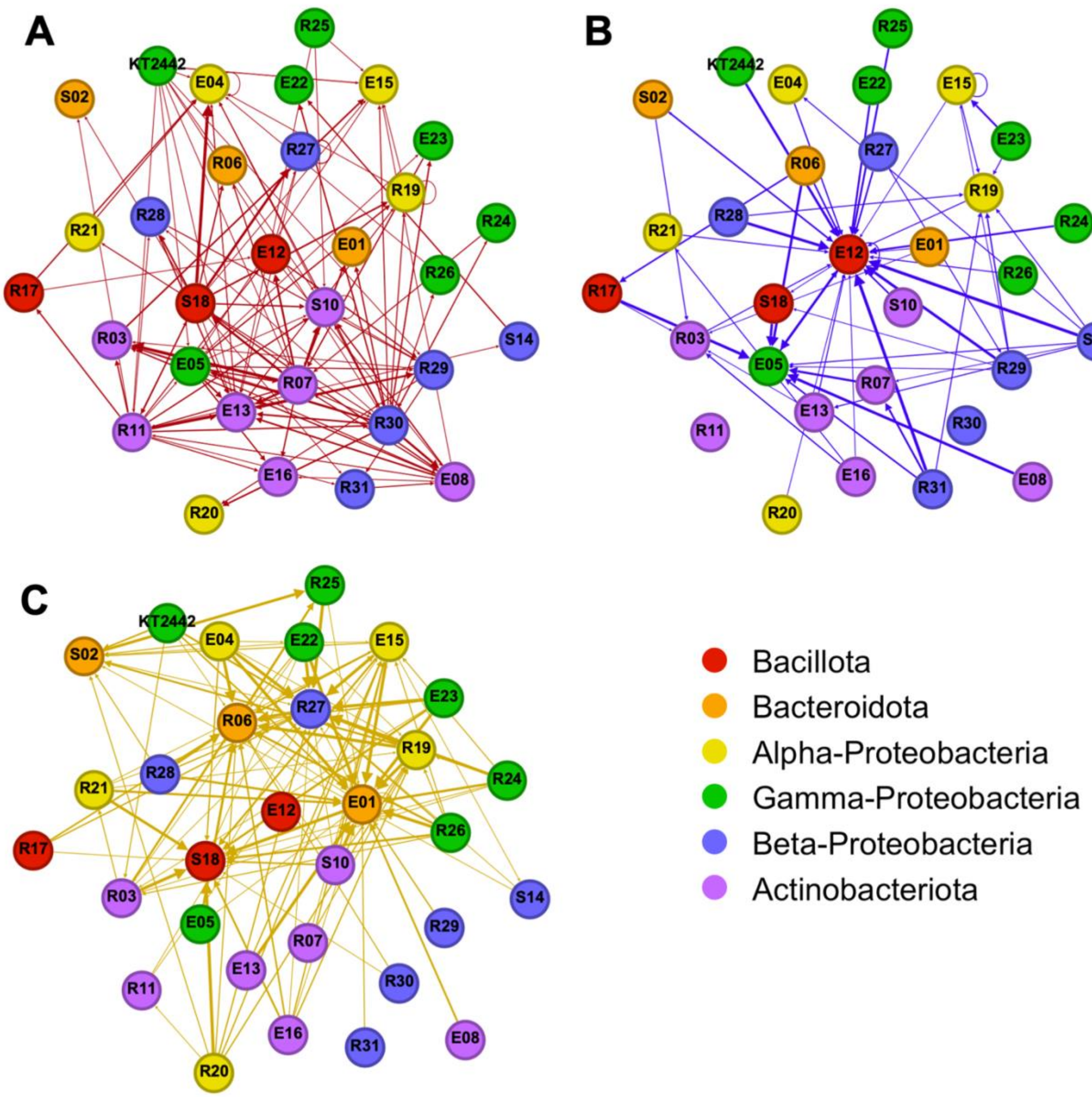


Figure S2. Interactions network for each MARSc member in the absence of exogenous Fe: (A) Inhibition of growth, (B) Stimulation of growth, and (C) Stimulation of colonial expansion/motility. Thickness of arrows indicates strength of the interaction. Only interactions greater than 1 mm in diameter are displayed.

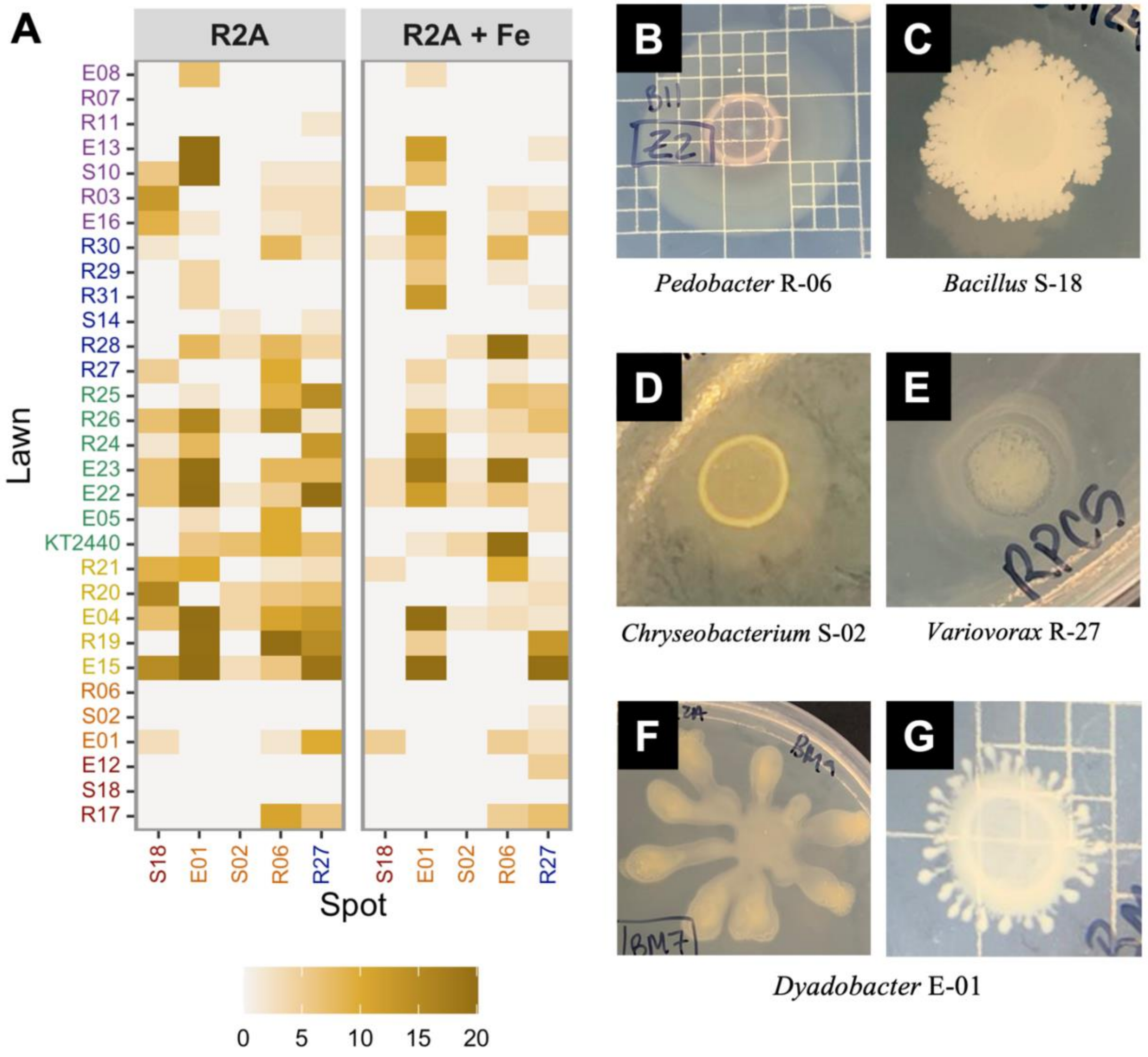


Figure S3. (A) Motility/colonial expansion of MARSc members (spots) during interactions with other strains (lawns), 96 hours post-inoculation. Motility/colony expansion was measured as millimeters of movement/expansion from the edge of the initial spot. Darker colors indicate greater colony expansion motility, with all measurements >20 mm reported as 20 mm. (B-G) Representative images showing the motility morphology of the 5 organisms who displayed colonial expansion.

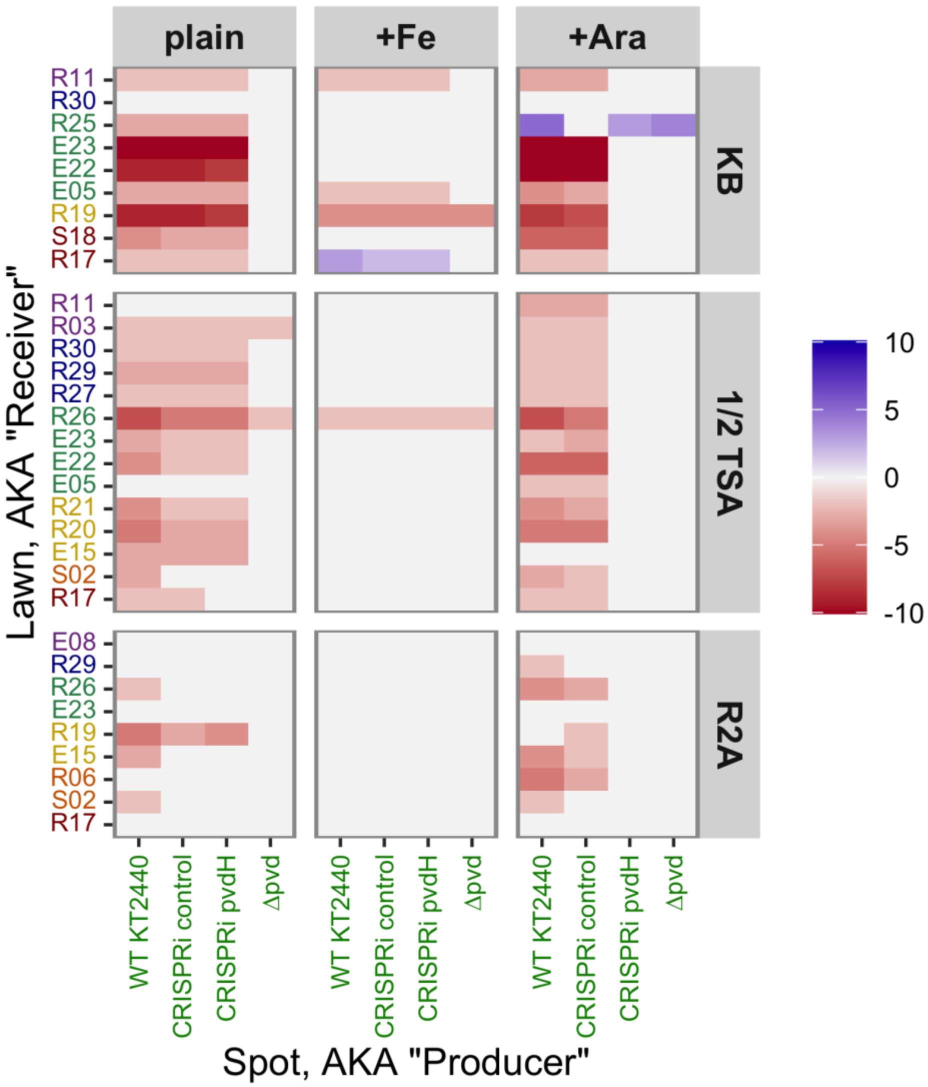


Figure S4. Pyoverdine production by *P. putida* KT2440 drives inhibitory interactions. KB, ½-TSA, and R2A media were either unamended, or amended with supplemental iron (50 µM $FeCl_3$ for KB, 35 µM for ½-TSA and R2A), or arabinose (67 mM). Arabinose was used to turn on the CRISPRi system, repressing *pvdH*. Blue indicates stimulation of growth, while red indicates inhibition of growth 2-3 days after inoculation. The darker the color, the more strongly the interaction was observed (measured in millimeters). Only MARSc members that showed decreased inhibition in the presence of exogenous iron are listed. See methods for description of strains used.

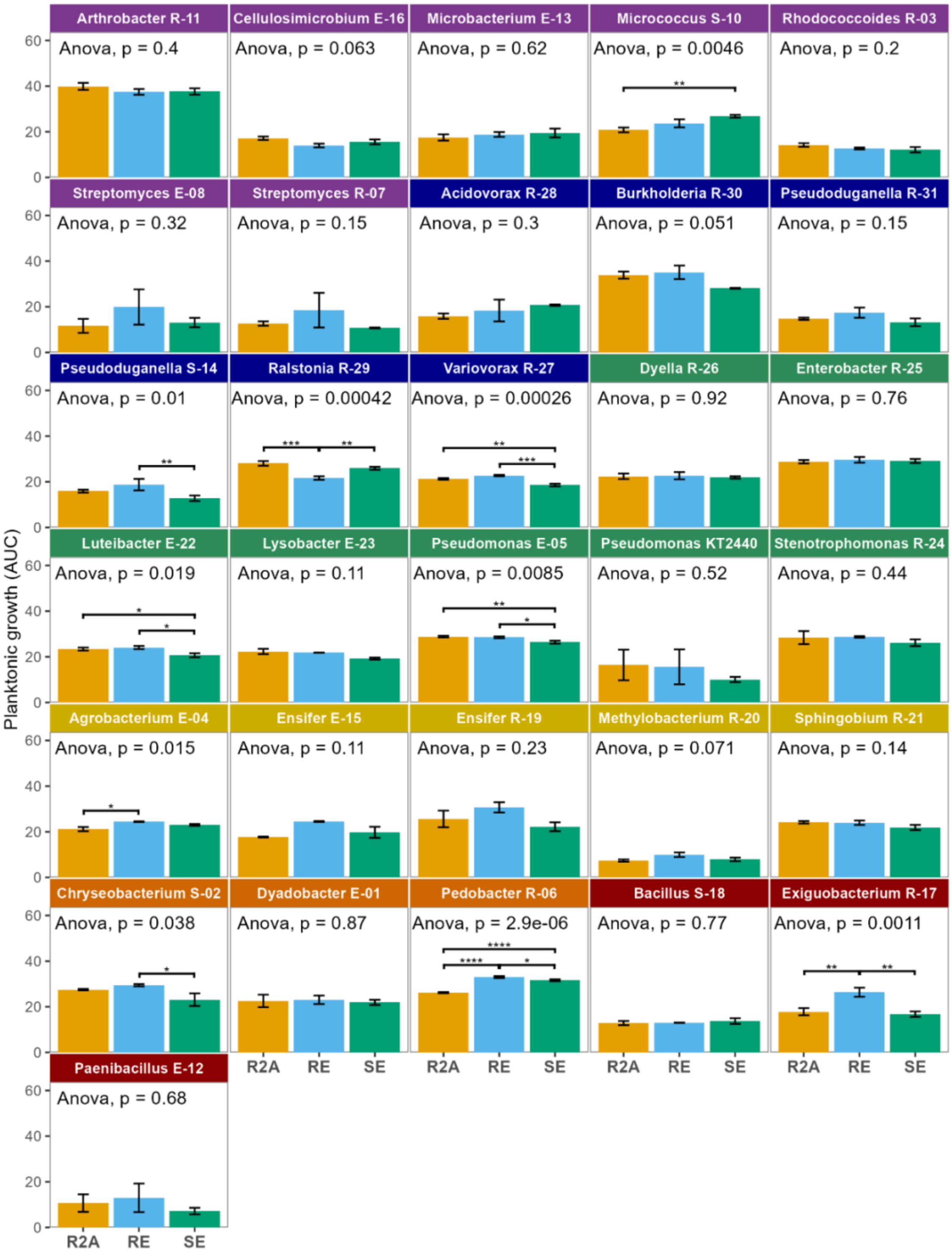


Figure S5 – Average Area Under the Curve (AUC) over a 44-hour growth period. Values are mean ± standard error. ANOVA p-values shown for each strain, with significant pairwise comparisons marked by bars and asterisks. Significance was determined with Tukey HSD. **** $p < 0.0001$, *** $0.0001 < p < 0.001$, ** 0.001 < p 0.01, * $0.01 < p < 0.05$

Table S1. Bacterial strains included in MARSc. Strain ID, genus and species, phylum and family, and NCBI genome accession numbers.

| **Strain ID** | **Genus/species** | **Phylum/family** | **Genome accession number** |
|---|---|---|---|
| E-12 | *Paenibacillus silvae* | Bacillota/Paenibacillaceae | CP183972 |
| R-17 | *Exiguobacterium sp* | Bacillota/Bacillaceae | CP184372-CP184373 |
| S-18 | *Bacillus velezensis* | Bacillota/Bacillaceae | CP184364 |
| E-01 | *Dyadobacter endophyticus* | Bacterioidota/Cytophagaceae | CP183969 |
| S-02 | *Chryseobacterium sp.* | Bacterioidota/Flavobacteriaceae | CP183989 |
| R-06 | *Pedobacter sp.* | Bacterioidota/Sphingobacteraceae | CP183977 |
| E-15 | *Ensifer adhaerens* | α-Proteobacteria/Rhizobiaceae | CP184365-CP184368 |
| R-19 | *Ensifer sp.* | α-Proteobacteria/Rhizobiaceae | CP184569-CP184571 |
| E-04 | *Agrobacterium radiobacter* | α-Proteobacteria/Rhizobiaceae | CP184251-CP184253 |
| R-20 | *Methylobacterium fujisawaense* | α-Proteobacteria/Methylobacteraceae | CP183980 |
| R-21 | *Sphingobium sp.* | α-Proteobacteria/Sphingomondaceae | CP184374-CP184376 |
| E-22 | *Luteibacter sp.* | γ-Proteobacteria/Xanthomonadaceae | CP183975 |
| E-23 | *Lysobacter soli* | γ-Proteobacteria/Xanthomonadaceae | CP183976 |
| R-24 | *Stenotrophomonas maltophilia* | γ-Proteobacteria/Xanthomonadaceae | CP183981 |
| E-05 | *Pseudomonas brassicacearum* | γ-Proteobacteria/Pseudomonadaceae | CP183970 |
| KT2440 | *Pseudomonas putida* | γ-Proteobacteria/Pseudomonadaceae | NC_002947.4 |
| R-25 | *Enterobacter ludwigii* | γ-Proteobacteria/Enterobacteriaceae | CP183982 |
| R-26 | *Dyella kyungheensis* | γ-Proteobacteria/Rhodanobacteraceae | CP183983 |
| R-27 | *Variovorax sp.* | β-Proteobacteria/Commonadaceae | CP183984 |
| R-28 | *Acidovorax facilis* | β-Proteobacteria/Commonadaceae | CP183985 |
| R-29 | *Ralstonia sp.* | β-Proteobacteria/Burkholderiaceae | CP184377-CP184380 |
| R-30 | *Burkholderia ambifaria* | β-Proteobacteria/Burkholderiaceae | CP184381-CP184383 |
| S-14 | *Pseudoduganella sp.* | β-Proteobacteria/Oxalobacteraceae | CP183991 |
| R-31 | *Pseudoduganella sp.* | β-Proteobacteria/Oxalobacteraceae | CP183986 |
| R-07 | *Streptomyces sp.* | Actinomycetota/Streptomycetaceae | CP183978 |
| E-08 | *Streptomyces sp.* | Actinomycetota/Streptomycetaceae | CP183971 |
| R-03 | *Rhodococcus fascians* | Actinomycetota/Nocardioidaceae | CP184369-CP184371 |
| S-10 | *Micrococcus yunnanesis* | Actinomycetota/Micrococcaceae | CP183990 |
| R-11 | *Arthrobacter sp.* | Actinomycetota/Micrococcaceae | CP183979 |
| E-13 | *Microbacterium sp.* | Actinomycetota/Microbacteriaceae | CP183973 |
| E-16 | *Cellulosimicrobium sp.* | Actinomycetota/Promicromonosporaceae | CP183974 |

Table S2. Traits of MARSc members. Biofilm formation in the presence/absence of MARSc metabolites. Production of various community- or plant-relevant compounds by MARSc members, including autoinducer-2 (AI-2) and production or degradation of indole acetic acid (IAA). Increase: biofilm formation in filter plates or spent media was greater than in normal well plates. Decrease: biofilm formation in filter plates or spent media was less than in normal well plates. *: There was no change to biofilm formation when in the filter plate, but there was a change in spent media. No increase/decrease: there was a change in biofilm formation when in the filter plate, which was not replicated by the spent media. YES: production of AI-2 was detected. Produce: Significant increase in IAA concentration of the culture. Degrade: Significant decrease in IAA concentration of the culture. x: no change to biofilm formation, no production of AI-2, or no production/degradation of IAA.

| | Impact of community metabolites on biofilm formation via… | | | | | |
|---|---|---|---|---|---|---|
| **Strain ID** | Living cells in filter plate (R2A) | Spent media (R2A) | Living cells in filter plate (R2A+RE) | Spent media (R2A+RE) | AI-2 production | IAA production/ degradation |
| *Arthrobacter* R-11 | x | x | x | x | x | Produce |
| *Micrococcus* S-10 | x | x | x | x | x | Produce |
| *Microbacterium* E-13 | x | x | x | x | x | x |
| *Cellulosimicrobium* E-16 | x | x | x | x | x | x |
| *Streptomyces* E-08 | x | x | x | x | x | x |
| *Streptomyces* R-07 | x | x | x | x | x | x |
| *Rhodococcus* R-03 | x | x | x | x | x | x |
| *Acidovorax* R-28 | x | x | x | x | x | x |
| *Variovorax* R-27 | x | x | x | x | x | Degrade |
| *Ralstonia* R-29 | x | x | Decrease | Decrease | x | x |
| *Burkholderia* R-30 | x | x | x | x | x | Degrade |
| *Pseudoduganella* R-31 | x | x | x | x | x | Degrade |
| *Pseudoduganella* S-14 | Increase | *No increase* | Increase | *No increase* | x | Degrade |
| *Pseudomonas* E-05 | Increase | *No increase* | Decrease | Decrease | x | x |
| *Pseudomonas putida* KT2440 | Increase | Increase | x | Decrease* | x | x |
| *Enterobacter* R-25 | Decrease | Decrease | x | x | YES | Produce |
| *Stenotrophomonas* R-24 | x | x | x | x | x | x |
| *Lysobacter* E-23 | x | x | Increase | *No increase* | x | x |
| *Luteibacter* E-22 | x | x | x | x | x | x |
| *Dyella* R-26 | x | x | x | x | x | x |
| *Sphingobium* R-21 | Decrease | Decrease | x | x | x | Produce |
| *Methylobacterium* R-20 | x | x | x | x | x | x |
| *Agrobacterium* E-04 | x | x | x | x | x | Produce |

| | | | | | | |
|---|---|---|---|---|---|---|
| *Ensifer* E-15 | x | x | x | x | x | x |
| *Ensifer* R-19 | x | x | x | x | x | Produce |
| *Chryseobacterium* S-02 | Decrease | *No decrease* | Decrease | *No decrease* | x | x |
| *Dyadobacter* E-01 | x | x | x | x | x | x |
| *Pedobacter* R-06 | x | x | x | x | x | x |
| *Paenibacillus* E-12 | x | x | x | x | x | x |
| *Bacillus* S-18 | Increase | *No increase* | Increase | Increase | YES | x |
| *Exiguobacterium* R-17 | x | x | x | x | YES | x |

Table S3. Extracellular protease activity of MARSc members. Casein hydrolysis in ½ TSA and R2A measured over the course of 5 days. Values are the mean zone of clearance of 3 biological replications. NT = Not Tested.

| | **Zone of Clearing (cm)** | | | | | |
|---|---|---|---|---|---|---|
| | **Day 1** | | **Day 2** | | **Day 5** | |
| **Strain ID** | **1/2 TSA** | **R2A** | **1/2 TSA** | **R2A** | **1/2 TSA** | **R2A** |
| *Arthrobacter* R-11 | 2.17 | 2.38 | 2.65 | 2.83 | 3.02 | 3.48 |
| *Micrococcus* S-10 | 2.40 | 2.34 | 3.04 | 2.73 | 4.01 | 3.69 |
| *Microbacterium* E-13 | 0.67 | 1.13 | 0.67 | 0.67 | 0.67 | 0.67 |
| *Cellulosimicrobium* E-16 | 1.84 | 1.00 | 2.68 | 2.46 | 3.38 | 3.13 |
| *Streptomyces* E-08 | 1.40 | 2.18 | 2.23 | 2.39 | 2.43 | 2.93 |
| *Streptomyces* R-07 | 2.42 | 2.50 | 2.47 | 2.47 | 3.07 | 3.46 |
| *Rhodococcus* R-03 | *NT* | *NT* | *NT* | *NT* | *NT* | *NT* |
| *Acidovorax* R-28 | *NT* | *NT* | *NT* | *NT* | *NT* | *NT* |
| *Variovorax* R-27 | 1.00 | 1.00 | 1.00 | 1.00 | 1.00 | 1.00 |
| *Ralstonia* R-29 | *NT* | *NT* | *NT* | *NT* | *NT* | *NT* |
| *Burkholderia* R-30 | 2.42 | 2.62 | 3.28 | 3.46 | 4.11 | 4.32 |
| *Pseudoduganella* R-31 | 0.67 | 1.56 | 0.67 | 2.44 | 1.81 | 2.73 |
| *Pseudoduganella* S-14 | 1.00 | 1.48 | 1.00 | 2.37 | 1.92 | 2.50 |
| *Pseudomonas* E-05 | 1.00 | 1.00 | 1.00 | 1.72 | 1.83 | 1.60 |
| *Pseudomonas putida* KT2440 | 1.88 | 1.00 | 1.94 | 1.00 | 1.00 | 1.00 |
| *Enterobacter* R-25 | 1.64 | 1.00 | 1.84 | 1.00 | 1.75 | 1.00 |
| *Stenotrophomonas* R-24 | 2.55 | 2.56 | 2.98 | 3.14 | 3.49 | 3.96 |
| *Lysobacter* E-23 | 0.67 | 1.67 | 2.36 | 2.82 | 3.32 | 3.21 |
| *Luteibacter* E-22 | 1.00 | 1.86 | 2.94 | 3.16 | 4.08 | 4.01 |
| *Dyella* R-26 | 0.67 | 0.33 | 0.67 | 0.33 | 1.44 | 1.27 |
| *Sphingobium* R-21 | 1.42 | 1.44 | 1.74 | 1.78 | 2.04 | 1.88 |
| *Methylobacterium* R-20 | 0.33 | 0.83 | 1.50 | 1.44 | *NT* | *NT* |
| *Agrobacterium* E-04 | 1.00 | 1.00 | 1.00 | 1.00 | 1.00 | 1.33 |
| *Ensifer* E-15 | 1.00 | 1.00 | 1.00 | 1.00 | 1.00 | 1.00 |
| *Ensifer* R-19 | 1.00 | 1.10 | 1.00 | 1.51 | 1.00 | 1.51 |
| *Chryseobacterium* S-02 | 2.19 | 2.42 | 2.59 | 2.94 | 3.11 | 3.17 |
| *Dyadobacter* E-01 | 1.00 | 1.47 | 1.52 | 1.88 | 1.00 | 1.00 |
| *Pedobacter* R-06 | *NT* | *NT* | *NT* | *NT* | *NT* | *NT* |
| *Paenibacillus* E-12 | 1.56 | 0.67 | 2.54 | 1.58 | 2.59 | 1.90 |
| *Bacillus* S-18 | 2.64 | 2.62 | 2.69 | 2.25 | 2.71 | 2.27 |
| *Exiguobacterium* R-17 | 3.17 | 2.94 | 3.69 | 3.70 | 4.37 | 3.99 |

Table S4. Plant growth promoting genes significantly correlated with rhizosphere abundance and their PLaBAse categories and functions. Corr. denotes correlation, either negative (-) or positive (+). P-values were calculated for each nitrogen level (60N – low, 120N – moderate, 180N – high) using Kendall correlations adjusted for false discovery rate. "# of MARSc members" is the number of MARSc members annotated with that gene.

| | | | | P-value | | | | |
|---|---|---|---|---|---|---|---|---|
| Gene | PLaBAse categories | Functions | Corr. | 60N | 120N | 180N | # of MARSc members | Strain ID of MARSc genomes containing the gene |
| phoU / phoY | Phosphate solubilization, heavy metal detoxification | Phosphate transport | - | 0.042 | 0.042 | 0.042 | 29 | E-04, E-05, E-08, E-12, E-13, E-15, E-16, E-22, E-23, KT2440, R-03, R-06, R-07, R-11, R-17, R-19, R-20, R-21, R-24, R-25, R-26, R-27, R-28, R-29, R-30, R-31, S-02, S-10, S-18 |
| cueR | Bacterial fitness, heavy metal detoxification | Multidrug transport - efflux pump | - | 0.017 | 0.007 | 0.007 | 23 | E-04, E-05, E-08, E-12, E-15, E-16, E-22, E-23, KT2440, R-03, R-07, R-19, R-20, R-24, R-25, R-26, R-27, R-28, R-29, R-30, R-31, S-02, S-18 |
| mcp / tlpC / tlpA / dcrA | Chemotaxis, bacterial fitness | Methyl-accepting chemotaxis proteins | - | 0.029 | 0.029 | 0.065 | 21 | E-04, E-05, E-12, E-15, E-22, E-23, KT2440, R-17, R-19, R-20, R-21, R-24, R-25, R-26, R-27, R-28, R-29, R-30, R-31, S-14, S-18 |
| clsA_B / ybhO / ywiE | Neutralizing abiotic stress, phospholipid production, cell envelope remodeling, quorum sensing | Cardiolipin biosynthesis | - | 0.024 | 0.010 | 0.024 | 27 | E-04, E-05, E-12, E-13, E-15, E-16, E-22, E-23, KT2440, R-03, R-11, R-17, R-19, R-20, R-21, R-24, R-25, R-26, R-27, R-28, R-29, R-30, R-31, S-02, S-10, S-14, S-18 |
| grxC | Neutralizing abiotic stress | Oxidative stress-chaperones | - | 0.029 | 0.029 | 0.065 | 24 | E-04, E-05, E-08, E-12, E-13, E-15, E-22, E-23, KT2440, R-03, R-19, R-20, R-21, R-24, R-25, R-26, R-27, R-28, R-29, R-30, R-31, S-10, S-14, S-18 |
| oxlT | Plant derived substrate usage | Formate and oxalate transport | - | 0.060 | 0.012 | 0.028 | 22 | E-04, E-05, E-08, E-12, E-13, E-15, E-23, KT2440, R-03, R-07, R-11, R-19, R-20, R-21, R-25, R-26, R-27, R-28, R-29, R-30, R-31, S-14 |
| ppx / ppx_gppA | Plant derived substrate usage, phosphate solubilization | Exopolyphosphatase and purine metabolism | + | 0.034 | 0.034 | 0.034 | 28 | E-01, E-04, E-05, E-08, E-12, E-15, E-16, E-22, E-23, KT2440, R-03, R-06, R-07, R-11, R-17, R-19, R-20, R-21, R-24, R-25, R-26, R-27, R-28, R-29, R-30, R-31, S-02, S-10 |
| TC_SSS/ yerK/ opuE | Plant derived substrate usage, neutralizing abiotic stress | Proline transport | + | 0.015 | 0.015 | 0.033 | 19 | E-01, E-05, E-08, E-22, E-23, KT2440, R-06, R-07, R-11, R-17, R-20, R-25, R-27, R-28, R-29, R-31, S-02, S-10, S-18 |

| | | | | | | | | |
|---|---|---|---|---|---|---|---|---|
| rhaM / rhaU / yiiL | Plant derived substrate usage | Rhamnose degradation | + | 0.007 | 0.017 | 0.007 | 18 | E-01, E-04, E-08, E-13, E-15, E-22, R-06, R-07, R-11, R-19, R-21, R-25, R-26, R-27, R-29, R-30, R-31, S-14 |
| iolW | Plant derived substrate usage | Opine metabolism and inositol derivative degradation | + | 0.010 | 0.010 | 0.010 | 22 | E-01, E-04, E-05, E-08, E-16, E-22, E-23, R-06, R-07, R-11, R-17, R-20, R-21, R-24, R-25, R-26, R-27, R-29, R-30, S-02, S-10, S-18 |
| Malz | Plant derived substrate usage, plant cell wall degradation, EPS production | Glycosidases / glycohydrolases / glucosidase | + | 0.007 | 0.007 | 0.007 | 16 | E-01, E-08, E-16, E-22, E-23, R-06, R-07, R-11, R-21, R-24, R-25, R-26, R-29, R-30, R-31, S-02 |
| ywaD | Plant derived substrate usage, plant cell wall degradation, EPS production | Peptide metabolism | + | 0.028 | 0.060 | 0.028 | 13 | E-01, E-22, E-23, R-06, R-21, R-24, R-26, R-27, R-31, S-02, S-10, S-14, S-18 |
| ydfG | Plant derived substrate usage, plant cell wall degradation, EPS production | Pyrimidine metabolism | + | 0.007 | 0.017 | 0.007 | 27 | E-01, E-04, E-05, E-08, E-13, E-15, E-16, E-22, E-23, KT2440, R-03, R-06, R-07, R-11, R-17, R-19, R-21, R-24, R-25, R-26, R-27, R-29, R-30, R-31, S-02, S-10, S-18 |
| bioF | Root colonization | Vitamin B7 / biotin biosynthesis | + | 0.017 | 0.042 | 0.042 | 24 | E-01, E-04, E-05, E-15, E-22, E-23, KT2440, R-03, R-06, R-07, R-11, R-19, R-20, R-21, R-24, R-25, R-26, R-27, R-28, R-29, R-30, R-31, S-02, S-18 |
| Hsd | Xenobiotics degradation | Cholesterol degradation | + | 0.029 | 0.029 | 0.029 | 11 | E-01, E-16, E-23, KT2440, R-06, R-07, R-11, R-20, R-21, R-24, S-10 |
| hyfR | Nitrogen acquisition | Hydrogenase biosynthesis | + | 0.037 | 0.037 | 0.037 | 4 | E-01, E-05, R-06, R-20 |
| qorB | Plant vitamin production | Vitamin K biosynthesis | + | 0.042 | 0.017 | 0.017 | 22 | E-01, E-04, E-05, E-08, E-12, E-15, E-16, E-22, R-03, R-06, R-07, R-11, R-17, R-19, R-25, R-26, R-27, R-29, R-30, R-31, S-02, S-14 |
| desB / desD / desA | Plant signal - linolenic acid production | Linolenic acid biosynthesis | + | 0.007 | 0.007 | 0.007 | 11 | E-01, E-08, E-13, E-23, R-03, R-06, R-11, R-24, R-26, R-29, S-02 |
| tnp | Bacterial fitness | Transposases | + | 0.037 | 0.037 | 0.037 | 15 | E-01, E-04, E-05, E-15, KT2440, R-03, R-06, R-07, R-19, R-20, R-21, R-24, R-29, R-30, S-10 |

345

Table S5. CRISPRi strains used in this study. $Rf^R$ = resistant to rifampicin.

| **Strain name in text** | **Genotype** | **Reference** |
| --- | --- | --- |
| KT2440 | wild type *Pseudomonas putida* KT2440 ($Rf^R$ variant) | (32) |
| CRISPRi control | KT2440::AraC/*pBAD*-dCas9 (pJMP1237) + pSpyB-Tn7sg | (2, 14) |
| CRISPRi *pvdH* | KT2440::AraC/*pBAD*-dCas9 (pJMP1237) + pSpyB-*pvdH*-77 | (2, 14) |
| Δ*pvd* | pyoverdine-deficient EZ-Tn5 transconjugant of KT2440 | this study |

### References


1. Brenner EA, Blanco M, Gardner C, Lübberstedt T. 2012. Genotypic and phenotypic characterization of isogenic doubled haploid exotic introgression lines in maize. Mol Breeding 30:1001–1016.

2. Roghair Stroud MN, Vang DX, Halverson LJ. 2024. Optimized CRISPR Interference System for Investigating Pseudomonas alloputida Genes Involved in Rhizosphere Microbiome Assembly. ACS Synth Biol 13:2912–2925.

3. Murashige T, Skoog F. 1962. A Revised Medium for Rapid Growth and Bio Assays with Tobacco Tissue Cultures. Physiol Plant 15:473–497.

4. Gest H, Favinger JL, Madigan MT. 1985. Exploitation of $N_2$ -fixation capacity for enrichment of anoxygenic photosynthetic bacteria in ecological studies. FEMS Microbiology Letters 31:317–322.

5. Davis AS, Hill JD, Chase CA, Johanns AM, Liebman M. 2012. Increasing Cropping System Diversity Balances Productivity, Profitability and Environmental Health. PLOS ONE 7:e47149.

6. Ma L, Shi Y, Siemianowski O, Yuan B, Egner TK, Mirnezami SV, Lind KR, Ganapathysubramanian B, Venditti V, Cademartiri L. 2019. Hydrogel-based transparent soils for root phenotyping in vivo. Proceedings of the National Academy of Sciences of the United States of America 166:11063–11068.

7. Paulsen AA, Vang DX, Halverson LJ. 2025. Complete genome sequences of 37 bacteria in a maize rhizosphere synthetic community. Microbiology Resource Announcements 0:e00496-25.

8. Bay G, Lee C, Chen C, Mahal NK, Castellano MJ, Hofmockel KS, Halverson LJ. 2021. Agricultural Management Affects the Active Rhizosphere Bacterial Community Composition and Nitrification. mSystems 6:10.1128/msystems.00651-21.

9. Olson RD, Assaf R, Brettin T, Conrad N, Cucinell C, Davis JJ, Dempsey DM, Dickerman A, Dietrich EM, Kenyon RW, Kuscuoglu M, Lefkowitz EJ, Lu J, Machi D, Macken C, Mao C, Niewiadomska A, Nguyen M, Olsen GJ, Overbeek JC, Parrello B, Parrello V, Porter JS, Pusch GD, Shukla M, Singh I, Stewart L, Tan G, Thomas C, VanOeffelen M, Vonstein V, Wallace ZS, Warren AS, Wattam AR, Xia F, Yoo H, Zhang Y, Zmasek CM, Scheuermann RH, Stevens RL. 2023. Introducing the Bacterial and Viral Bioinformatics Resource Center (BV-BRC): a resource combining PATRIC, IRD and ViPR. Nucleic Acids Res 51:D678–D689.

10. Lozano GL, Bravo JI, Garavito Diago MF, Park HB, Hurley A, Peterson SB, Stabb EV, Crawford JM, Broderick NA, Handelsman J. 2019. Introducing THOR, a model microbiome for genetic dissection of community behavior. mBio 10:1–14.

11. Vetsigian K, Jajoo R, Kishony R. 2011. Structure and evolution of streptomyces interaction networks in soil and in silico. PLoS Biology 9:e1001184.

12. Shannon P, Markiel A, Ozier O, Baliga NS, Wang JT, Ramage D, Amin N, Schwikowski B, Ideker T. 2003. Cytoscape: A Software Environment for Integrated Models of Biomolecular Interaction Networks. Genome Res 13:2498–2504.

13. Bastian M, Heymann S, Jacomy M. 2009. Gephi: An Open Source Software for Exploring and Manipulating Networks. ICWSM 3:361–362.

14. Peters JM, Koo BM, Patino R, Heussler GE, Hearne CC, Qu J, Inclan YF, Hawkins JS, Lu CHS, Silvis MR, Harden MM, Osadnik H, Peters JE, Engel JN, Dutton RJ, Grossman AD, Gross CA, Rosenberg OS. 2019. Enabling genetic analysis of diverse bacteria with Mobile-CRISPRi. Nature Microbiology 4:244–250.

15. Ceri H, Olson ME, Stremick C, Read RR, Morck D, Buret A. 1999. The Calgary Biofilm Device: New Technology for Rapid Determination of Antibiotic Susceptibilities of Bacterial Biofilms. J Clin Microbiol 37:1771–1776.

16. Chodkowski JL, Shade A. 2017. A Synthetic Community System for Probing Microbial Interactions Driven by Exometabolites. mSystems 2:4–7.

17. Chodkowski JL, Shade A. 2024. Bioactive exometabolites drive maintenance competition in simple bacterial communities. mSystems 9:e00064-24.

18. Thoenen L, Giroud C, Kreuzer M, Waelchli J, Gfeller V, Deslandes-Hérold G, Mateo P, Robert CAM, Ahrens CH, Rubio-Somoza I, Bruggmann R, Erb M, Schlaeppi K. 2023. Bacterial tolerance to host-exuded specialized metabolites structures the maize root microbiome. Proceedings of the National Academy of Sciences 120:e2310134120.

19. Custer GF, Dibner RR. 2020. Modified Methods for Loading of High-Throughput DNA Extraction Plates Reduce Potential for Contamination. JoVE 61405.

20. Srivathsan A, Lee L, Katoh K, Hartop E, Kutty SN, Wong J, Yeo D, Meier R. 2021. ONTbarcoder and MinION barcodes aid biodiversity discovery and identification by everyone, for everyone. BMC Biology 19:217.

21. Paulsen AA, LaSarre B, Delp D, Beattie GA, Halverson LJ. 2026. A Bioinformatic Pipeline for Consensus Taxonomic Classification of Long-Read Amplicons. bioRxiv https://doi.org/10.64898/2026.04.29.721641.

22. Liu C, Cui Y, Li X, Yao M. 2021. microeco: an R package for data mining in microbial community ecology. FEMS Microbiology Ecology 97:fiaa255.

23. Fernandes AD, Reid JN, Macklaim JM, McMurrough TA, Edgell DR, Gloor GB. 2014. Unifying the analysis of high-throughput sequencing datasets: characterizing RNA-seq, 16S rRNA gene sequencing and selective growth experiments by compositional data analysis. Microbiome 2:15.

24. Fernandes AD, Macklaim JM, Linn TG, Reid G, Gloor GB. 2013. ANOVA-Like Differential Expression (ALDEx) Analysis for Mixed Population RNA-Seq. PLOS ONE 8:e67019.

25. Schwengers O, Jelonek L, Dieckmann MA, Beyvers S, Blom J, Goesmann A. 2021. Bakta: rapid and standardized annotation of bacterial genomes via alignment-free sequence identification. Microbial Genomics 7:000685.

26. Sequeira JC, Rocha M, Alves MM, Salvador AF. 2022. UPIMAPI, reCOGnizer and KEGGCharter: Bioinformatics tools for functional annotation and visualization of (meta)-omics datasets. Computational and Structural Biotechnology Journal 20:1798–1810.

27. Patz S, Gautam A, Becker M, Ruppel S, Rodríguez-Palenzuela P, Huson D. 2021. PLaBAse: A comprehensive web resource for analyzing the plant growth-promoting potential of plant-associated bacteria. bioRxiv https://doi.org/10.1101/2021.12.13.472471.

28. Alterman JL, Vang DX, Roghair Stroud MN, Halverson LJ, Kraus GA. 2020. Ozonolysis of Alkynes - A Flexible Route to Alpha-Diketones: Synthesis of AI-2. Organic Letters 22:7424–7426.

29. Taga ME, Xavier KB. 2011. Methods for Analysis of Bacterial Autoinducer-2 Production. Current Protocols in Microbiology 23:1C.1.1-1C.1.15.

30. Gordon SA, Weber RP. 1951. Colorimetric Estimation of indoleacetic Acid. Plant Physiol 26:192–195.

31. Gu Z. 2022. Complex heatmap visualization. iMeta 1:e43.

32. Espinosa-Urgel M, Ramos J-L. 2004. Cell Density-Dependent Gene Contributes to Efficient Seed Colonization by *Pseudomonas putida* KT2440. Appl Environ Microbiol 70:5190–5198.
